# Local Ancestry Inference for Complex Population Histories

**DOI:** 10.1101/2023.03.06.529121

**Authors:** Alice Pearson, Richard Durbin

## Abstract

It has become apparent from ancient DNA analysis, that the history of many human populations from across the globe are often complex, involving multiple population split, admixture, migration and isolation events. Local ancestry inference (LAI) aims to identify from which ancestral population chromosomal segments in admixed individuals are inherited. However, ancestry in existing LAI tools is characterised by a discrete population identity, a definition which is limited in the context of a complex demographic history involving multiple admixture events at different times. Moreover, many LAI tools rely on a reference panel of present day genomes that act as proxies for the ancestral populations. For ancient admixture events, these proxy genomes are likely only distantly related to the true ancestral populations. Here we present a new method that leverages advances in ancient DNA sequencing and genealogical inference to address these in issues in LAI. The method applies machine learning to tree sequences inferred for ancient and present day genomes and is based on a deterministic model of population structure, within which we introduce the concept of path ancestry. We show that the method is robust to a variety of demographic scenarios, generalises over model misspecification and that it outperforms a leading local ancestry inference tool. We further describe a downstream method to estimate the time since admixture for individuals with painted chromosomes. We apply the method to a large ancient DNA dataset covering Europe and West Eurasia and show that the inferred admixture ages are a better metric than sample ages alone for understanding movements of people across Europe in the past.

## Introduction

Despite a wealth of data, patterns in modern genomes alone are difficult to interpret as they are an indirect measure of past events. Moreover, much of the genetic variation which was present in past populations does not exist in the modern gene pool due to ancient demographic events that exacerbate drift, such as bottlenecks and isolation. Ancient DNA provides a genetic snapshot of a time before these processes have taken place, meaning we regain some of the lost information. As a result, the use of aDNA has transformed our understanding of human origins and evolution in recent years [1] showing that the histories of human populations are often complex, involving periods of isolation and migrations with admixture events [2].

Admixture events occur when there is migration between divergent populations and they interbreed. Chromosomes of the resulting admixed individuals will originate in one of the two ancestral populations. With each generation there is recombination, meaning that over time chromosomes in the admixed population will be a mosaic of chunks originating in the two ancestral populations. Local ancestry inference (LAI) is the process of decomposing admixed chromosomes into these ancestral chunks and assigning each chunk an ancestry label. Many tools are available that perform LAI due to its importance for understanding population structure, migration history and disease risks [3, 4].

Early LAI methods [5, 6] use unlinked markers that are characteristic of populations, so called ancestry informative markers (AIMs). The hidden Markov model (HMM) structure employed by these methods relied on the assumption that markers are independent and so do not model background LD but admixture LD alone. With the decrease in cost of genotyping in recent years, the ability to deconvolve local ancestry with greater resolution using denser SNP data became possible but violated the assumption of independence between SNPs. Methods that leverage this denser data emerged [7, 8] that employed a pruning step which makes sure SNPs are unlinked in the ancestral populations. However, not explicitly modelling background LD within ancestral populations prevents the use of many informative SNPs that are linked. Later methods now both utilise denser SNP data and model background LD together with admixture LD by using extensions of the basic HMM approach and leveraging haplotype frequencies, which vary more between populations than SNP frequencies [9–11]. More recently machine learning has been used for local ancestry inference and has shown to be computational efficient and accurate. Furthermore, the ever increasing availability of genomic data provides training data for supervised methods [12, 13].

Complex population histories pose a problem for LAI as most of the existing methods require sequences that represent a discrete ancestral populations to form a reference panel [7, 9, 10, 12, 13]. In reality, haplotypes are inherited from generation to generation, often through a complex demographic history, possibly involving multiple admixture events between many populations. An ancestral population may itself be an admixed population from an earlier event in history. In other words, populations are more like a braided river than a sequence of well-defined discrete population identities. Thus, assignment of a haplotype in an admixed chromosome to a single ancestral population in a reference panel does not inform us of ancestry further back in time or take into account the genealogical relationship of that ancestral population to other reference populations. Some LAI tools allow multiple reference populations that could represent many populations involved in a focal population history from different time periods [7, 9, 12, 13], but haplotypes in focal admixed samples can only be assigned to one of these ‘ancestries’ from the reference panel when in fact two or more assignments may be true. Moreover, the reference panel is often made up of present day samples that are closely related proxies of the true ancestral populations and thus the accuracy of ancestry assignment varies with how well the reference populations represent the true ancestry populations. The relationship of present day genomes to ancestral populations that existed deeper in time becomes more distant making LAI less accurate for more ancient admixture events.

Here we develop a local ancestry inference method that takes the genealogical relationships of ancestral populations into account and leverages the power of ancient samples. For this we redefine ‘ancestry’ as no longer a discrete population identity, but a complete path back in time through the population history. The path that a haplotype takes backwards in time from a focal individual is fully informative about its local ancestry. Ancient samples are used to represent ancestral populations along these paths as they are more closely related to the true ancestral groups involved in the admixture events. The method involves building tree sequences, a representation of the ancestral recombination graph, between a large set of sample sequences using RELATE [14]. RELATE enables sample sequences to be placed in the past [15] and so we can incorporate ancient samples into the tree sequences [15]. We then train a neural network, using a simulated training set, that classifies the ancestry path for each sample haplotype in a genomic region given the local tree and a known population history.

There has been extensive research on the population history of Europeans [2, 16, 17] and the broad picture is well established. Therefore, we used a large dataset of present day and high quality shotgun sequenced, imputed ancient genomes (MesoNeo dataset) to test the method. We constructed a model in msprime [18] that represents the major ancestry flows contributing to modern European genomes over the last 50,000 years, from which we simulated tree sequence and variation data. Local ancestry inference performed by a neural network, trained on simulated data from this model, is as good if not better than inference using GNOMix [13], a leading LAI tool. Additionally, we showed the method is robust to a range of simulated demographic scenarios and model misspecification.

Lastly, we applied a technique to infer the time since admixture of admixed MesoNeo individuals using the rate of switching between painted segments and thus gain insight into the patterns of admixture event that occurred in history between major ancient European groups.

## Method and Materials

### Concept of path ancestry

We explain in more detail the issue current LAI tools face when using a reference panel of proxy ancestral population from which only one population can represent the ancestry of each admixed haplotype. Figure 1 shows a diagrammatic example of a population structure with two consecutive admixture events and two population split events; population *C* is an ancestral population to population *E.* A haplotype in *E* can be assigned to both C and *A;* both are true at the same time. Existing LAI methods could include both C and A as discrete ancestral reference populations, producing confusing results or choose either C or A for the reference panel and only use half the available data. In other words, all LAI methods treat ancestral populations as discrete entities with no genealogical relationships with each other, requiring them to effectively draw a line in the past time and take the populations existing at that time as ‘pure’ ancestral populations.

**Fig. 1.**
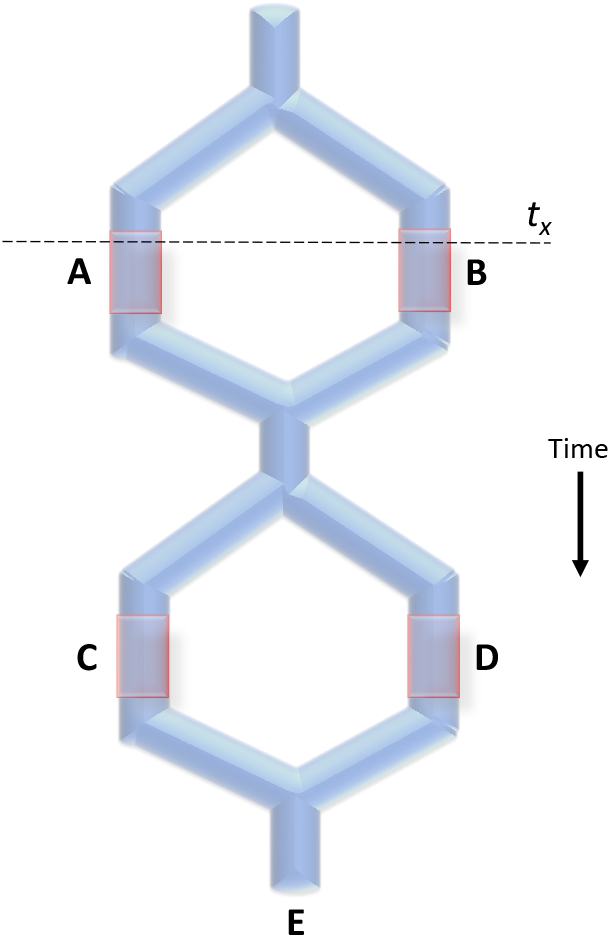
Population structure with paths. Schematic of a structured meta-population that goes through two population split events and two admixture events. Populations marked *A, B,C* and *D* are ‘ancestral populations’ and population *E* is the admixed population whose local ancestry is of interest.

The concept of ‘path’ ancestry takes time and genealogical relationships of ancestral populations into account. We redefine ‘ancestry’, not as a discrete population identity, but a complete path back in time through the population history. The path that a haplotype takes backwards in time from a focal individual is fully informative about its local ancestry: By determining what populations a haplotype has been carried by inheritance through a structured population history, the relationship of the haplotype to all relevant historical and admixing populations is established.

Returning to the example in Figure 1; take population E as the population whose haplotypes we wish to perform LAI on. We have representative samples from populations *A, B, C*, and *D* from within this structure which are termed ‘ancestral populations’. Between these four ancestral populations, there are four paths that haplotypes could have taken backwards in time from population *E*, through multiple populations:

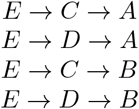

By using paths as local ancestry labels instead of discrete population identities, we are able to use all available data from all four ancestral populations and assign four different labels that convey meaningful information about the history of a haplotype.

Ancient samples within this framework are a huge advantage: The further back in time a population existed, the more difficult it is to find a closely related proxy population that exists today and from whom genomic data is available. Ancient samples are likely to be less diverged from the true ancestral populations and so act as better representatives that existed before one or many admixture and split events.

### Constructing a path model of European population history

Using the extensive amount of previous work elucidating the genetic history of Europeans, we put forward a standardised model of European population structure. Fig 2 shows a schematic of this demographic model that describes the population structure in Europe during the last 50,000 years. Shortly after the expansion of anatomically modern humans into Eurasia, there is a population split 15,000 years ago (1500 generations ago) between the Northern Europeans (NE), who continued moving northwest into Europe, and West Asians (WA) who stayed more locally in the Levant and South Caucasus area. The WA population then splits to form the Caucasus hunter gatherers/early Iranians (CHG) and the Anatolians (Ana) ~24,000 years ago (800 generations ago). Within the NE, ~18,000 years ago (600 generations ago), the western hunter gatherers (WHG) and eastern hunter gatherers (EHG) diverge. At that point, the four separate populations that make up present day European ancestry are distinct in the model. The timing of divergence between the four distinct groups appears to correspond to the onset of last glacial maximum [17, 19], during which time these population became isolated from each other in refugia.

**Fig. 2.**
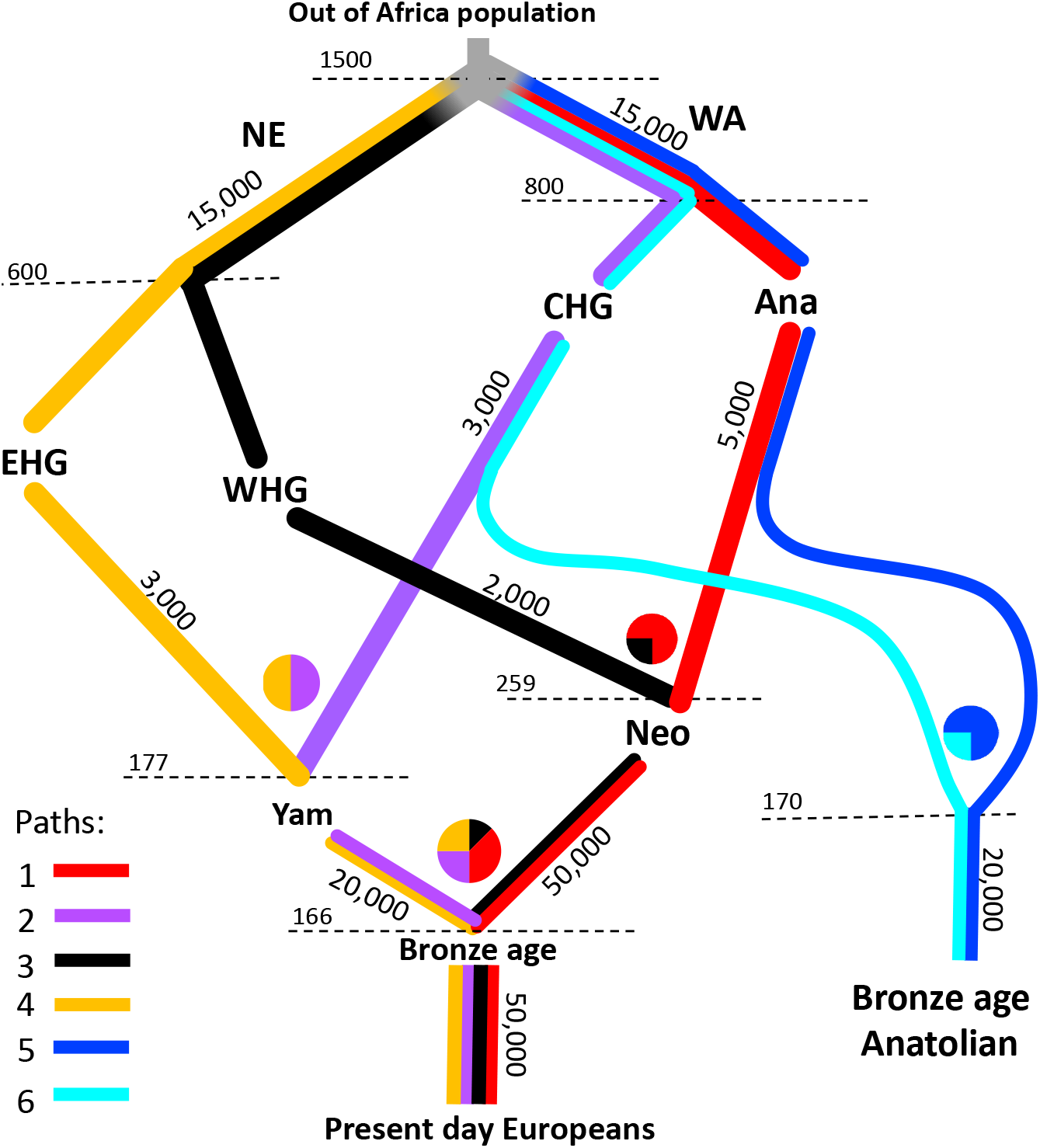
Schematic diagram of the model of European population structure. Population labels are abbreviated. Bronze Age = Bronze Age European population, Yam = Yamnaya steppe, Neo = Neolithic farmers, WHG = Western hunter gatherers, CHG = Caucasus hunter gatherers, EHG = Eastern hunter gatherers, Ana = Anatolian farmers, NE = Ancient Northern Europeans, WA = Ancient West Asians. Effective population sizes are shown along edges and admixture fractions are displayed as pie charts. The timing of population splits and admixture events are shown at the dotted lines in units of generations ago.

The model subsequently describes how admixture between these populations leads to the modern European gene pool. At the start of the Neolithic around 9,000 years ago, farmers from Anatolian moved into Europe. These incoming Anatolian farmers admixed with the local WHG ~7,800 years ago (259 generations ago) to form the Neolithic farmer population [20, 21] and their ancestry reaches Britain by 6,000 years ago [22].

During the late Neolithic, steppe populations, characterised by the Yamnaya (Yam) culture, appear as a mix of EHG and CHG/Iranian ancestries [17, 19, 23]“-5,300 years ago (177 generations ago). At a similar time, the Bronze Age Anatolian group (BAA) represent a later Anatolian population formed from admixture of CHGs with Ana ~5,100 years ago (170 generations ago). Around the start of the Bronze Age ~4,900 years ago (166 generations ago), migrants with Yam ancestry moved west into Europe and had a profound impact on the genetic landscape by admixing with the Neolithic farmers to form the European Bronze Age population, leading to present day Europeans [16, 20, 23–25].

Present day Europeans are often described as a three-way mix of WHG, Anatolian Farmer and steppe Yamnaya [2]. However, the Yamnaya ancestry itself resolves into EHG and CHG ancestry [17, 26] and these two groups have different deep origins as seen in Fig 2 [17]. Therefore, we propose a model of four ancestral streams that lead to present-day Europeans.

Six different paths that haplotypes can take from any sampled individual are shown in different colours in Fig 2. Path 1 = red, starts from the present day Europeans, going back through Neolithic farmers, Anatolian farmers, West Asians to the root. Path 2 = purple, starts at present day Europeans, going back through the Yamnaya, Caucasus hunter gatherers, then West Asians to the root. Path 3 = black, starts with present day Europeans, going back through Neolithic farmers to Western hunter gatherers and then through Northern Europeans to the root. Path 4 = orange, starts at present day Europeans, going back through the Yamnaya to Eastern hunter gatherers and then through Northern Europeans to the root. Paths 5 = blue, starting in the Bronze Age Anatolians and joins path 1 part. Path 6 = cyan, starts in Bronze Age Anatolians and joins path 2. When paths overlap, lineages from all overlapping paths can coalesce.

We constructed our model of European population history in msprime [18], a coalescent simulator. The population sizes and admixture fractions are shown in the schematic in Fig 2. This model has been submitted to the stdpopsim catalogue (https://github.com/popsim-consortium/stdpopsim/blob/main/stdpopsim/catalog/HomSap/demographic_models.py) and is publicly available for users to simulate data from.

### Comparison of simulated data to MesoNeo genomes

The MesoNeo dataset is a large collection of both ancient and present day samples. While the dataset contains samples from all over the world, most are from people that lived in the last 10,000 years in Eurasia. The dataset is composed of 1,490 genomes that have been imputed to 3.7 million SNPs once filtered for coverage (>0.1X), low imputation quality, close relatives and >0.5 imputation INFO score (See [27] for data availability).

To test how well our model represents the true history of Europeans, we compared data simulated from the model to the MesoNeo genomes. Based on PCA (Supplementary Figure S11) we subset the 1,492 genomes from the MesoNeo dataset to those that were tightly clustered in the groups relevant to the genetic history of Europeans, so that only samples diagnostic of each population in our model were kept. This totalled 476 diploid individuals (952 haploid) including 91 GBR 1000 Genomes samples. During simulation, samples were taken from populations and times to match each real diploid sample (Table **??**), using their radiocarbon or context dates. We aimed to create a simulated dataset that closely matched our MesoNeo subset genetically.

Upon simulation, data for sample individuals is produced, in the form of both VCF files and tree sequence files. We simulated chromosomes of length 200 Mbp with a recombination rate of 1e-8 per bp per generation and neutral mutations dropped onto the branches at a rate of 1.25e-8 per bp per generation. It is also possible to simulate using a real human chromosome recombination map in which case the sequence length is given by the file and a variable recombination rate across the sequence in applied. To manipulate and examine trees in tree sequence files we use the tskit python API.

We performed Principal Component Analysis on the simulated genotype data to compare the structure to that of the real data. Both the MesoNeo chromosomes and nine simulated chromosomes were filtered for minor allele frequency ¿5%. We used EIGENSOFT smartpca [28] with the outlier removal option disabled to perform PCA with no projection on both the imputed MesoNeo samples together 1000 Genomes GBR samples and the simulated samples.

In Figure 3 the cluster for each simulated ancient population falls in the same vicinity of PCA space as the real MesoNeo samples and the variation explained by PC1 and PC2 is comparable. Overall, the similarity of the PCA suggests that a lot of the underlying structure that determines how these groups relate to each other is captured by the demographic model.

**Fig. 3.**
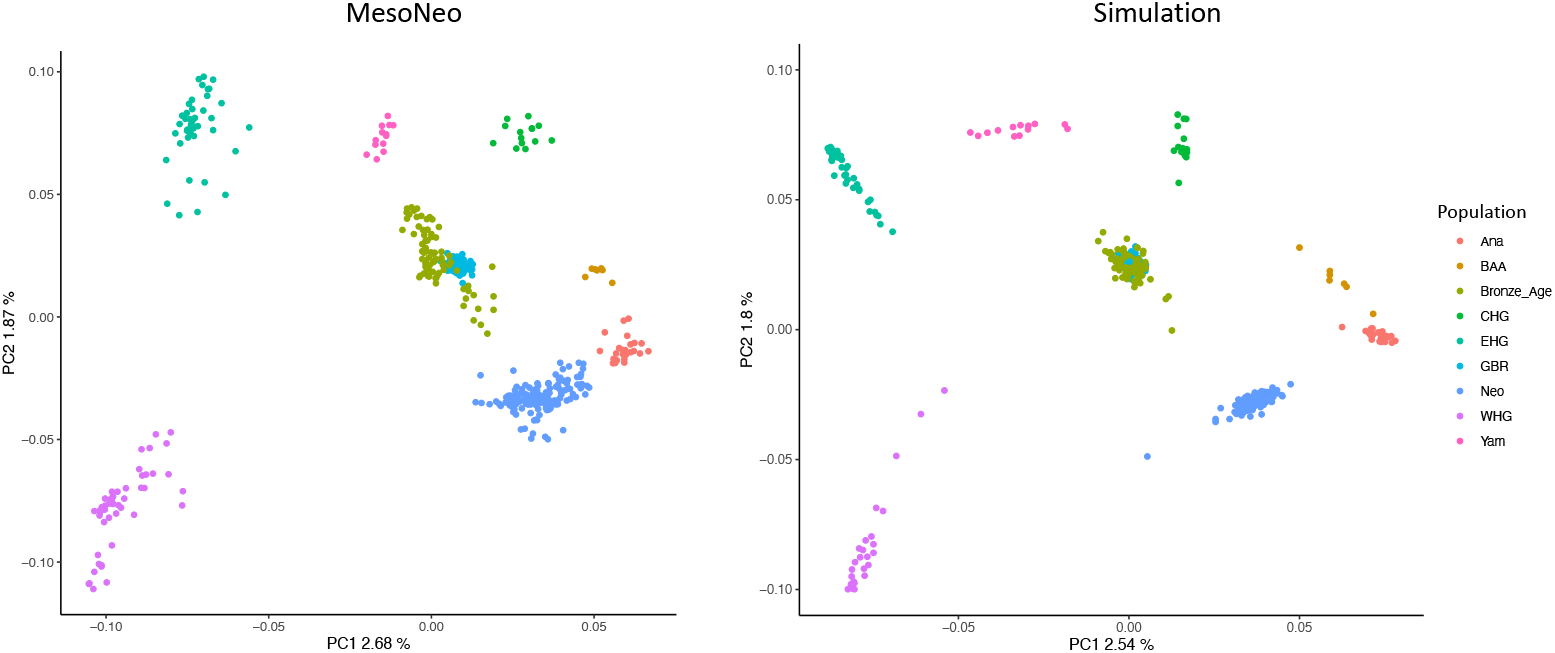
Comparison of simulated and MesoNeo PCA.

Weir and Cockerham weighted F_st_ statistics were calculated over all chromosomes using VCFtools between all pairwise populations. The same was done for the nine simulated sequences. The results shown in Figure 4 display the F_st_ pairwise values in the simulated data plotted against those in the real data. The correlation of F_st_ values between the real and simulated data is appreciable at 0.96. This means the demographic model produces data with relative population divergences that are very similar to that in the real data, suggesting the model represents the real population structure well.

**Fig. 4.**
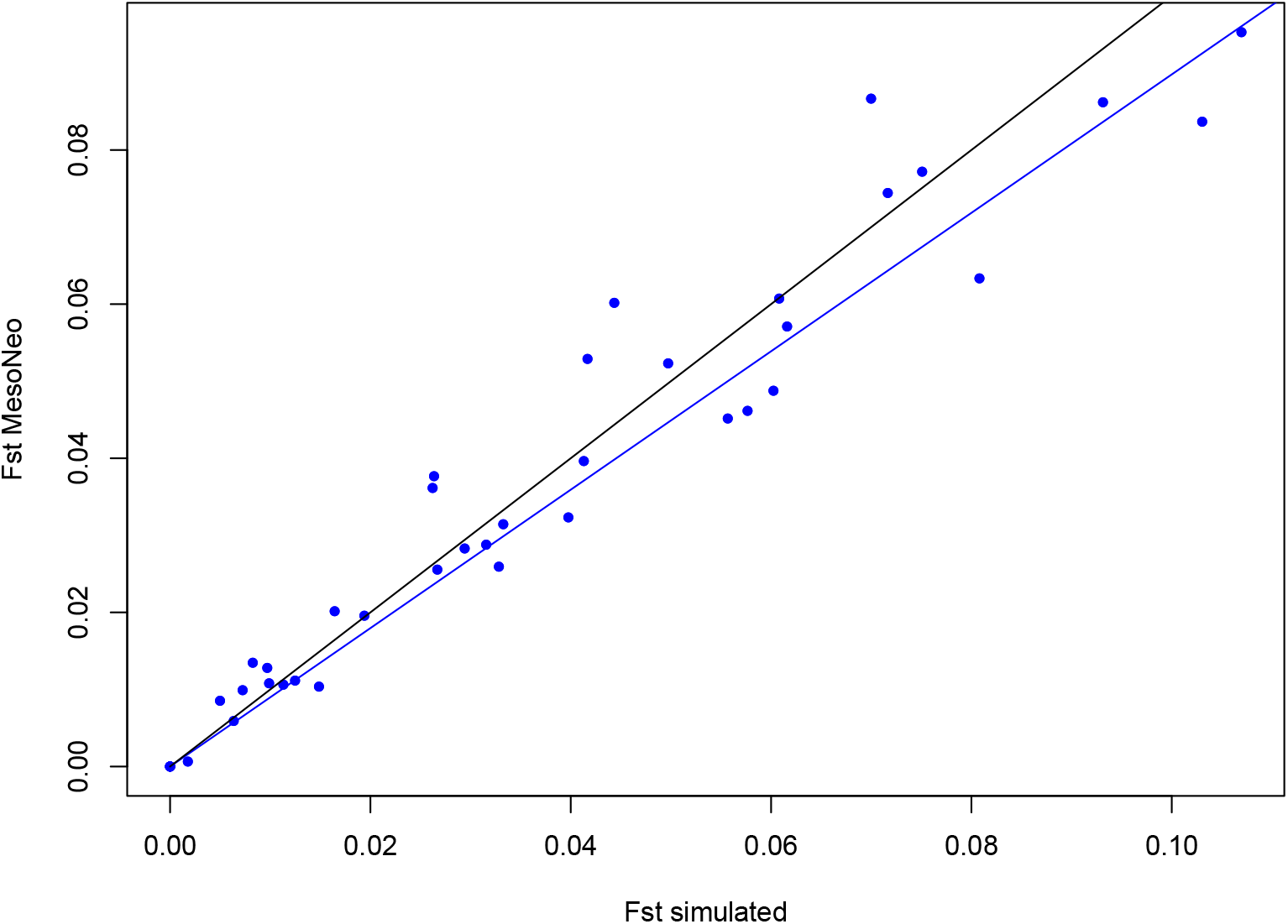
Pairwise F_st_ correlation.

The relationships of these ancient groups and present day Europeans to each other and the split/admixture times are well established and needed no adjustments in the model. However, to produce PCA and F_st_ results in simulated that matched the real data, the population sizes shown in Fig 2 needed some tuning. Both the F_st_ analysis and PCA final results suggest that, while the model is undoubtedly not entirely correct, it produces simulated data that looks close enough to the true data, by these measurements, that we assume the model represents the real population history of Europeans.

### A method for estimating local path ancestry

Our method infers the path that segments of a chromosome have taken through a population history. Each segment is covered by a single marginal tree in a tree sequence. Figure 5 depicts how the path ancestry of focal haplotype can be determined from the tree relating that haplotype to all other sample haplotypes: We traverse up the tree from the focal haplotye, jumping to successive parent nodes towards the root i.e parent, grandparent, great-grandparent etc. If we know the population identity of each of the internal nodes traversed to along the way, then we know the path. This information is recorded in tree seqeunces simulated by msprime and so the path ancestry of haplotypes in simulated tree sequences is straight-forward to find with tskit (https://tskit.dev/tskit/docs/stable/introduction.html).

**Fig. 5.**
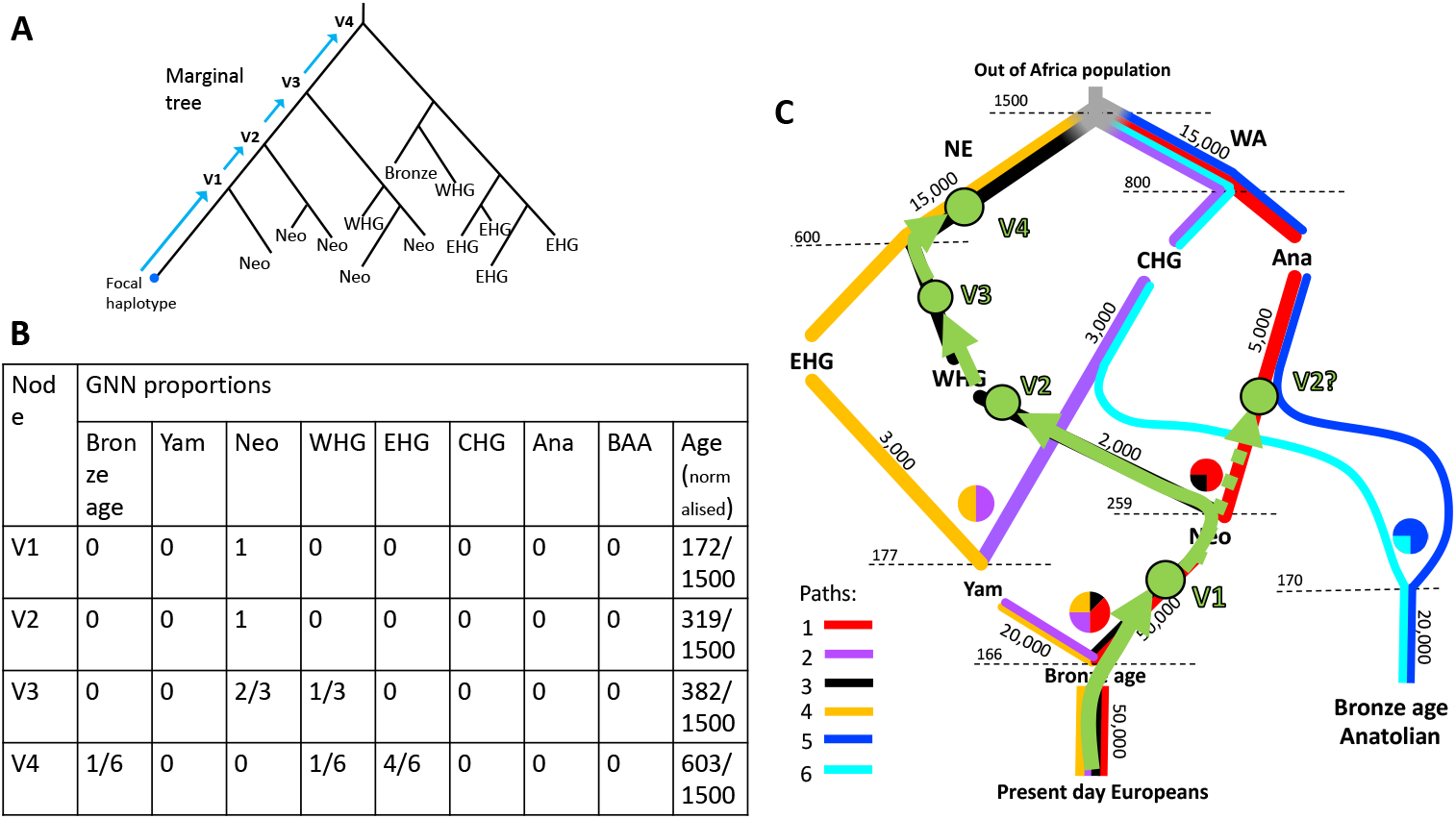
Overview of GNN extraction for a European population. (A) shows an example marginal tree that relates the focal haplotype to all other haplotypes in the dataset. Samples are shown by their population labels. (B) shows the GNN matrix determined from the marginal tree, traversing up four nodes towards the root V1-V4. (C) shows how the path can be determined within the model of population history given the GNN matrix by mapping the nodes to the paths.

However in RELATE tree sequences, inferred from simulated or real variant data, the population label of internal nodes is not known. We therefore adapted the concept of Genealogical Nearest Neighbours (GNNs) [29] to identify the population identity of each internal node encountered during traversal from a focal admixed haplotype. This involves recording the proportion of each ancestral group that makes up the sample leaves below each internal node, not including leaves seen at the previous nodes in traversal. The ordered collection of all *x* GNN distributions of all *x* nodes examined during a tree traversal reflects the path that the focal haplotype has taken to the root. Figure 5 demonstrates how these ordered GNNs can be used to determine the path.

We need to be able to assign paths to GNNs from millions of sample haplotypes so we implemented supervised machine learning, specifically a convolutional neural network. First we simulate variant and tree sequence data from a demographic model of the history of the populations under analysis, such as our model of European population structure. Next we infer RELATE tree sequences from the simulated genotype data. We then extract training GNNs from the RELATE trees, and train a neural network to predict the path given the true path label extracted from the corresponding simulated tree sequences. By training on GNNs extracted from RELATE trees, we aim to ‘train in’ any bias from the RELATE inference so that when we apply the neural network to tree sequences inferred by RELATE from the real data, these biasses are somewhat accounted for.

Training data for the network is the set of GNN distributions for the first five informative nodes traversed towards the root extracted from the RELATE inferred trees. The GNN distributions are configured as a 5 x *n* matrix, one row per 5 informative nodes examined. Columns 1 - n contain the proportions, between 0 and 1, of leaves belonging to each of n ancient sampled groups and column n +1 contains the age of the node. Informative nodes are those that have at least one leaf from the set of ancient sampled groups. If the root of the tree is reached in less than five informative nodes, then the remaining rows of the matrix are filled with −15 as ‘padding’. The training labels signify the path extracted from the corresponding simulated tree sequences (determined simply using tskit).

The method also works when traversing up to the root and finding the path for ancient sample haplotypes. Therefore, we also train over GNNs extracted from admixed samples, while samples that are diagnostic of one path (e.g Figure 2 CHG, WHG, EHG and Ana) can be assigned by their population identity alone.

The method performs well for a range of demographic scenarios including variable number of paths and extent of population differentiation, as well as being robust to misspecification in the underlying model (Supplementary notes S2, S3).

### Training and testing a neural network on simulated European genomes

We simulated three 200Mbp tree sequences from our model of European population structure, using different random seeds. RELATE tree sequences were inferred from the corresponding simulated VCF files. Default parameters for RELATE were used; 1.25e-8 mutation rate and starting population size estimate of haplotypes of 30,000.

We extracted 10,000 training pairs of GNNs and true labels from each of the five admixed sample populations (GBR, Bronze Age, Yamnaya, Neolithic farmers and Bronze Age anatolians), 50,000 pairs in total. GNNs were taken from the trees covering evenly spaced sites across three tree sequences output from RELATE, avoiding most of the correlation between trees. True path labels were taken from the trees covering the same sites in the corresponding simulated tree sequences. We trained the classifier to predict the path labels from the GNNs.

For testing, we generated five more simulated and RELATE inferred pairs of tree sequences and tested the classifier on 10,000 GNNs from each to obtain a mean accuracy of 93.12% with a standard deviation of 0.29% across the five tree sequences.

To test the precision of the classifier at identifying each path in each population, we pooled the testing GNNs from all five sequences and applied the classifier to each admixed population separately. Figure 6 shows the confusion matrices for each population, normalised by the sum of the rows to show the precision i.e of the labels assigned a class, what proportion are true positives. The GBR obtains the lowest accuracy. GBR samples have the greatest number of generations between sampling time and admixture time. More generations since the admixture event means more recombination events have broken down admixture LD, resulting in shorter tracts of ancestry. RELATE uses flanking SNPs to calculate the distance matrix and allows some relaxation on the mapping of SNPs when constructing a new tree along the chromosome. These properties make it more difficult to determine the switch of ancestry at the edges of tracts as there is some inertia in the changing of tree topologies compared to the true tree sequences. Shorter tracts produce more edges and therefore less accuracy in the GNN assignment. However, high accuracy is still maintained for all populations, even the GBR.

**Fig 6.**
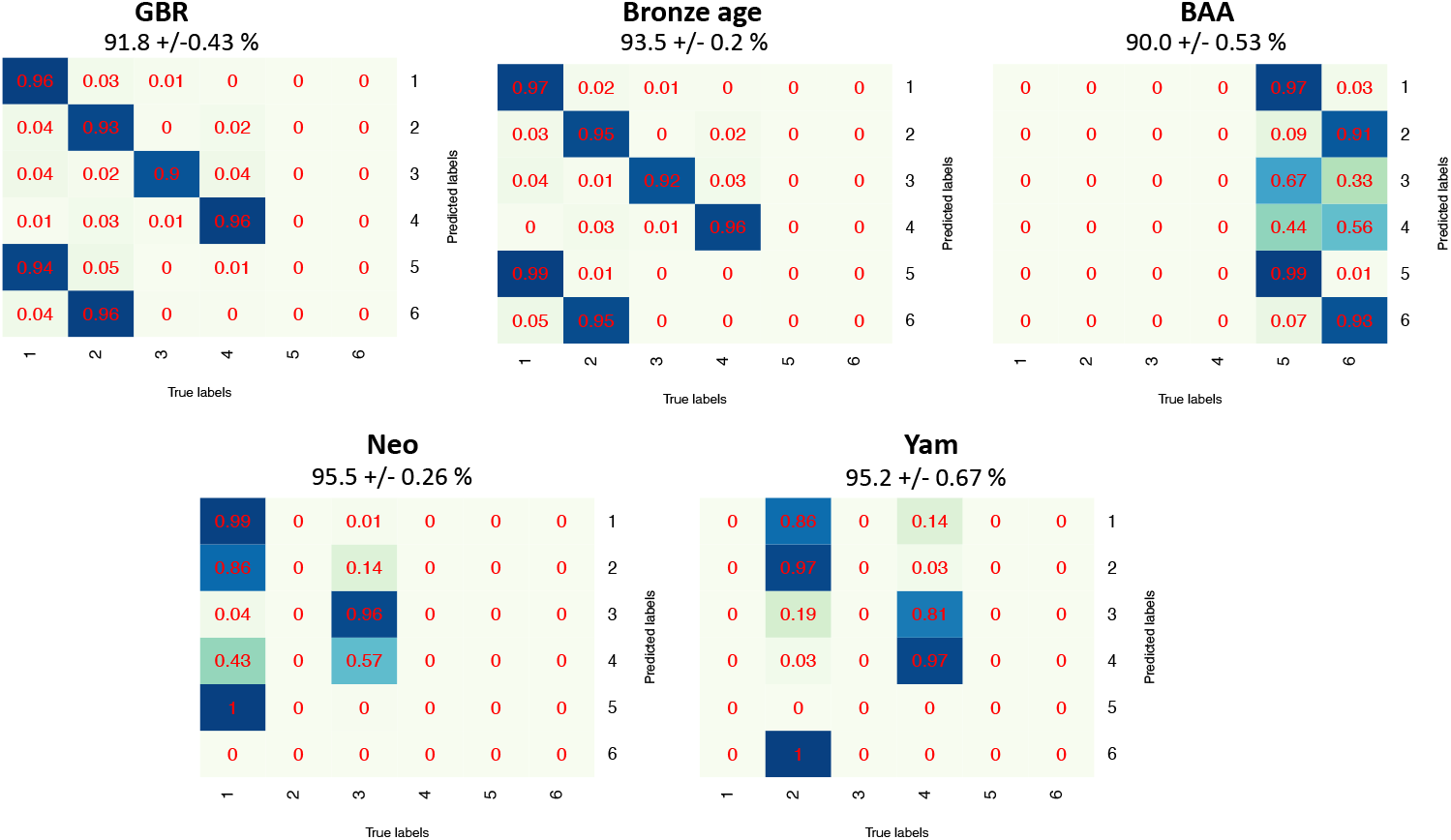
Confusion matrices per population when testing on simulated data. Values are normalised by the sum of rows to show the precision values, when testing a classifier trained on the model of European population structure.

Supplementary Figure S12 shows a classified painted chromosome alongside the true simulated chromosome. Noise appears as short tracts of correlated trees, within larger chunks of ancestry covering many sites in admixture LD.

### Comparison to an advanced LAI tool

GNOMix is a recent LAI tool that has been shown to outperform previous methods on whole genome data [13]. Like other LAI methods, GNOMix views each reference population as a discrete ancestral population with no awareness of the relationships between reference populations. It therefore is a good tool to compare our method against on simulated data of Europeans.

For the GNOMix reference panel we use samples from the four ‘path’ populations (EHG, WHG, CHG and Ana) as ancestral population that correspond to our paths 1 to 4 from the model. These are the populations that lie on one path only (Fig 2). For brevity we only test the four admixed populations as query sequences that are characterised by these four paths, GBR, Bronze Age, Neo and Yam, and do not test the BAA. It should be noted however that the BAA could be tested with GNOMix using the Ana and CHG as ancestral populations which would correspond to our paths 5 and 6.

We simulated five 200Mbp length sequences from the model of European population structure. We extracted ancestry predictions, produced by our method and by GNOMix, from evenly spaced sites across the five sequences. Inference with GNOMix was done using the default logistic regression base and xgboost smoother modules. All other parameters were default including a window size of 0.2cM for Bronze Age, Neo and Yamnaya samples. Because the GBR are 166 generations since admixture we decreased the window size to 0.02cM to account for the smaller admixture tracts. The mean and standard deviation for each admixed population, across the five sequences, obtained by each method were used to perform two-sample T-tests to test for a significant difference in accuracy (Table 1).

**Table 1.**
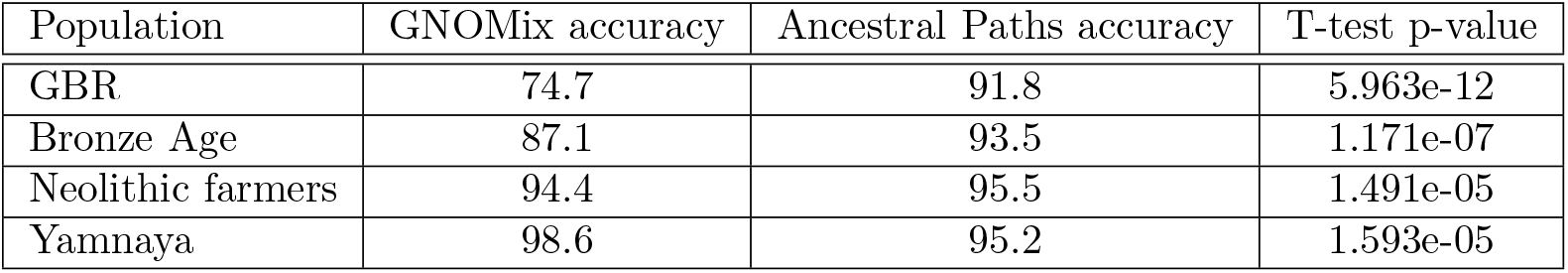
Comparison of accuracy to GNOMix for each of four admixed populations.

| Population | GNOMix accuracy | Ancestral Paths accuracy | T-test p-value |
| --- | --- | --- | --- |
| GBR | 74.7 | 91.8 | 5.963e-12 |
| Bronze Age | 87.1 | 93.5 | 1.171e-07 |
| Neolithic farmers | 94.4 | 95.5 | 1.491e-05 |
| Yamnaya | 98.6 | 95.2 | 1.593e-05 |

For the GBR, Bronze Age and Neo the our method is significantly more accurate. GNOMix was significantly more accurate for the Yam population. While GNOMix has high accuracies for populations closer to the time of admixture, classification of the present-day GBR samples is much less accurate than our method. Despite reducing the window size for GBR classification, it appears GNOMix struggles with smaller sized ancestry tracts. Our method displays high accuracy for all populations, demonstrating its versatility compared to GNOMix.

## Results

We inferred a RELATE tree sequence for each autosome of our subset of 476 MesoNeo genomes, using the fine scale recombination map for each chromosome, a mutation rate of 1.25e-8, and starting population size estimate of 30,000. All chromosomes for all samples were painted by the classifier trained on the model of European population structure.

### Estimation of time since admixture in Europe

We devised a technique to infer the time since admixture of individuals using the painted chromosomes. The method involves fitting exponential decay curves to the probability of being in the same path at two positions separated by varying genetic distances along a chromosome. The time since admixture in generations is extracted from a parameter of the exponential decay function (Supplementary note S4). This process is performed on all autosomes, treating homologous chromosomes independently. We obtain two estimates for each diploid sample, one estimate per ancestry, which are then combined as a weighted average to get an estimate of time since admixture per individual. For samples which consist of multiple paths, we combine the paths that make up each parent population involved in an admixture event, creating larger admixture chunks. For example, the Bronze Age admixture event is estimated from Bronze Age samples by combining paths 2 with 4 and paths 1 with 3 to represent the Yamnaya and Neolithic farmers admixing respectively.

We estimated the time of admixture for each sample in the three admixed MesoNeo populations; Neolithic farmers, Steppe Yamnaya and the Bronze Age population. The number of genomes over which this was performed was increased from our original 476 to include samples that fall in between groups in the PCA (Supplementary Figure S11), to explore whether these samples were the result of more recent admixture. The new total number of samples was 963, including all European 1000 Genomes samples. We showed that there was little decrease in accuracy when painting extra samples if we use the same set of 476 diagnostic samples in the GNNs (Supplementary note S3.5).

We calculated a date of admixture as the sample age plus the estimated time since admixture for each individual, combining the standard error of the radiocarbon age estimate and the time since admixture estimate. Given the coordinates of the archaeological sites, we fit spatiotemporal models of how time of admixture depends on latitude and longitude and compared these to the same models constructed using sample age alone.

#### Neolithic farmers

Of the 176 Neolithic farmers individuals in our subset dataset, we were able to estimate the time of admixture for 173. Figure 7 shows the inferred time of admixture in years against sample age. A linear regression to the points fits a model coefficient where every year younger in sample age increases the time since admixture in that sample by 0.28 years (p-value = 2.37e-06). This best fit line has a shallow gradient compared to the theoretical best fit line in which the Neolithic farmers are formed by one instantaneous admixture event happening 7,800 years ago. This suggests the migration of Neolithic farmers and admixture events between Neolithic farmers and WHG was less punctuated and more of a continuous process, ranging between 8,000 and 4,000 years ago.

**Fig 7.**
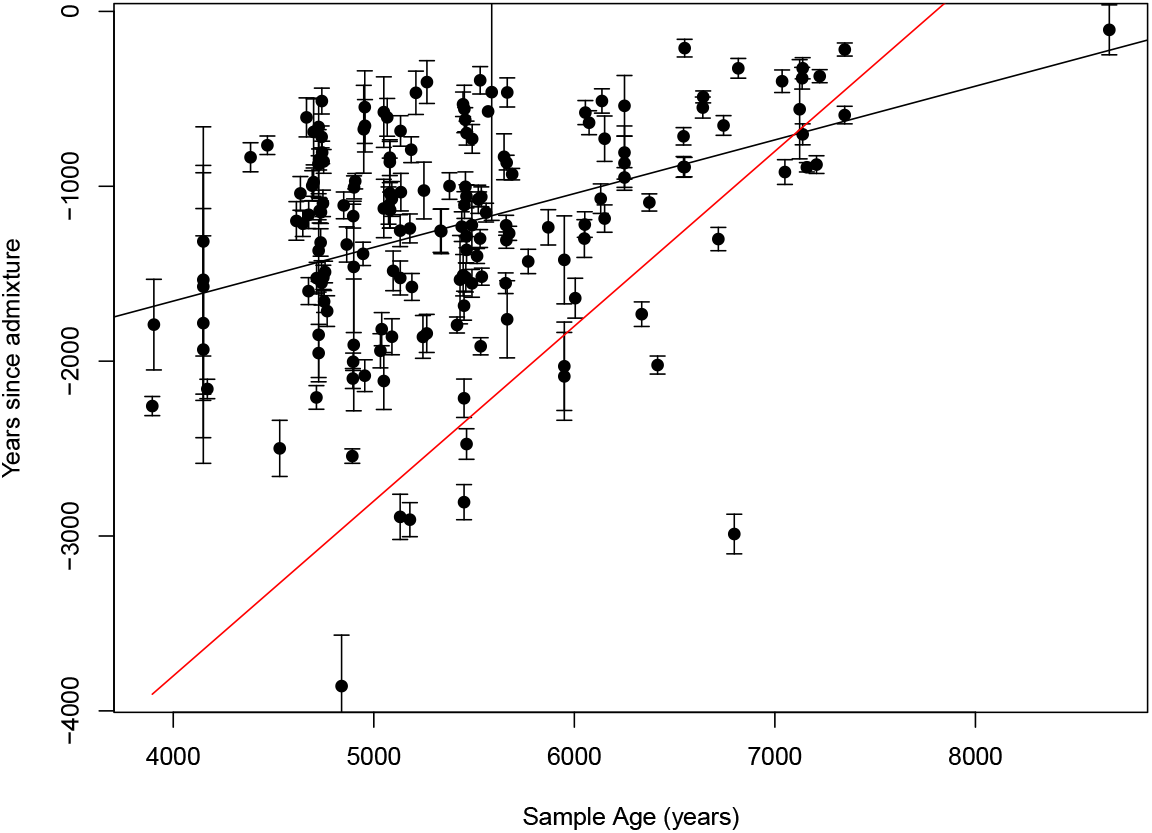
Correlation of years since admixture and sample age in 173 Neolithic farmer samples. The best fit line is coloured black. The theoretical best fit line, in the scenario of instantaneous admixture happening at 7,800 years ago, is coloured red.

Two theories exist for the route that Anatolian farmer ancestry took from its origin in Anatolian into the European continent. One is a path along the northern Mediterranean coast to Iberia first before expanding north and the second is an inland route following the Danube River, north and west.

We fit two linear models of Inferred admixture time ~ longitude + latitude and Sample age ~ longitude + latitude (Supplementary Table S8) which showed longitude and latitude are both highly significant predictors of admixture date. The model has a significant overall p-value of 5.947e-11 and explains 23.4% of the variance in the data. The predicted coefficients of longitude and latitude suggests that the admixture events occurs 25 years later with every degree east and 42 years later with every degree north. Similar results are shown in Figure 8 for which we performed linear interpolation of admixture time between the points of inferred admixture time of individual samples by kriging across Europe. The pattern of admixture of Anatolian farmers with WHG with time starts in Iberia and moves north east across Europe, implying the coastal route.

**Fig 8.**
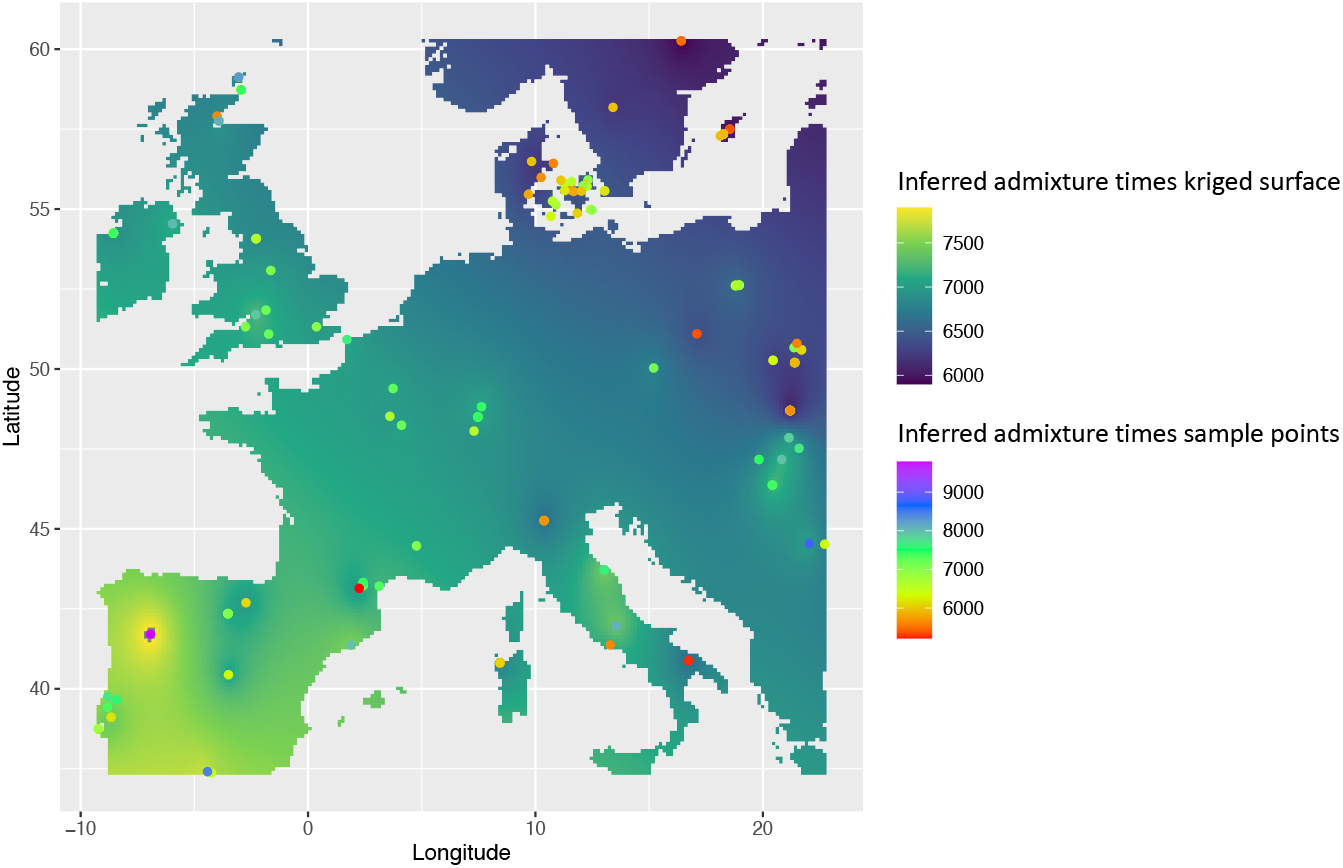
Linear interpolation of admixture time of Neolithic farmers across Europe. Map of Europe with points to show the archaeological position of Neolithic farmer samples coloured by the inferred admixture time in years before present. The surface colour is admixture time in years before present created by linear interpolation between inferred sample points.

The MesoNeo dataset contains few samples of Neolithic farmers from along the Mediterranean coast east of Spain, due to low sample availability, or perhaps poor preservation, and so we may not be capturing the earliest admixed samples. Alternatively, the first migrants moving along the coast may not have admixed with local hunter gatherer populations until they reached Iberia. Both scenarios would be consistent with our results and in the future, more samples from critical regions could help provide clarity.

In contrast, while longitude and latitude are also significant predictors of sample ages (Supplementary Table S8), the variance explained by such a model is only 8.4%. The model suggests a more gentle south east to north west gradient of older to younger sample ages. Overall, a model involving longitude and latitude to predict inferred admixture time is more informative than one predicting sample age, in terms of understanding the impact of Anatolian farmers in Europe.

#### Bronze Age population

The migration of the Yamnaya associated ancestry into Europe is typically thought to have been a fast migration, possibly accompanied by violence [30] and substantial alteration the environment [31].

Figure 9 shows the inferred time since admixture in years of the Yamnaya (path 2 and 4) and Neolithic farmers (paths 1 and 3) plotted against sample age. Of the 105 Bronze Age individuals in our MesoNeo subset, exponential curves could be fit to 97. A linear regression to these points implies that every year younger in sample archaeological age, the time since admixture in that sample increases by 0.78 years (p-value = 1.536e-11). A coefficient value of 1 would mean contemporaneous admixture over the whole continent, as shown by the theoretical best fit line in red. Such a high coefficient found for the Bronze Age admixture is a consistent with a rapid migration of the Yamnaya across the continent between 4000 and 5500 years ago, with admixture events quickly following.

**Fig 9.**
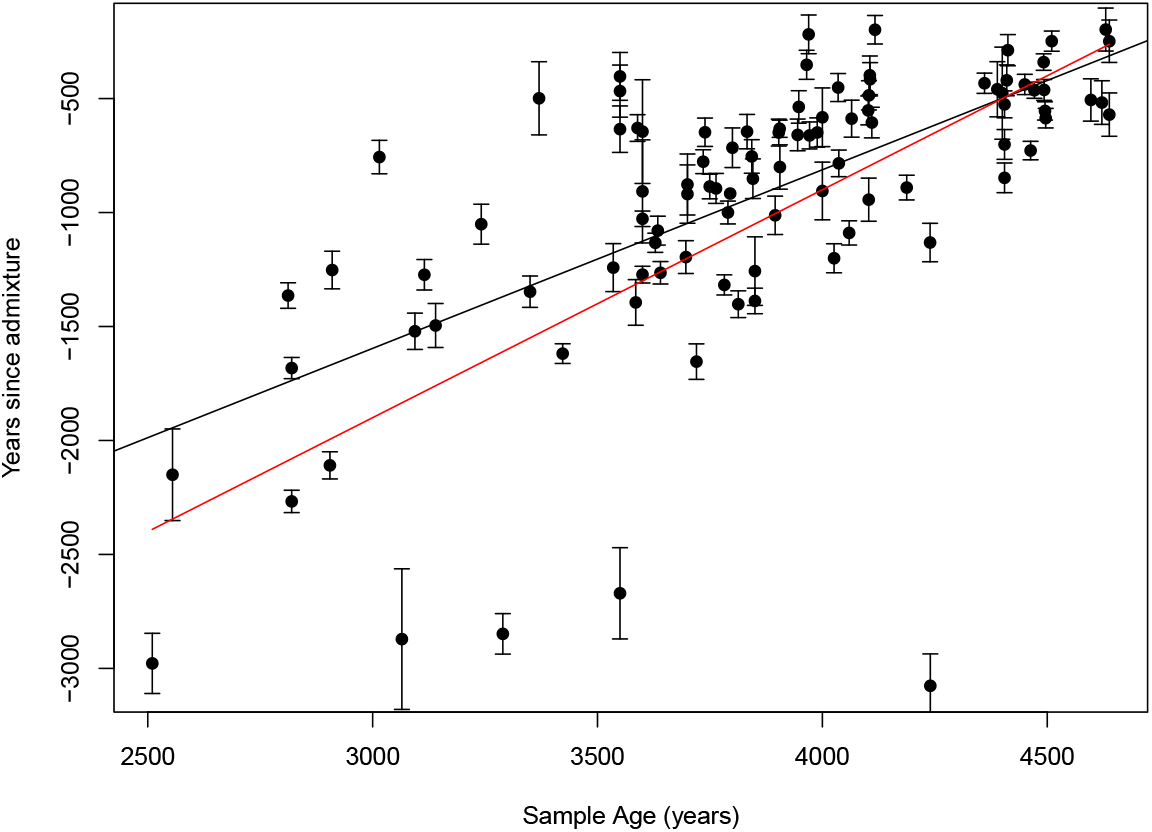
Correlation of years since admixture and sample age in 97 Bronze Age samples. The best fit line is coloured black. The theoretical best fit line, in the scenario of instantaneous admixture happening at 4,900 years ago, is coloured red.

Moreover, linear models incorporating longitude and latitude do not find either as a significant coefficients (Supplementary Table S9). This is supportive evidence of rapid movement of Yamnaya into Europe from the steppe with admixture following soon after the migration started, and then the admixture continuing with admixed individuals, producing no detectable variance in inferred admixture time with geography. The variance in the data explained by a model of longitude and latitude alone is small at only 4.9%. While this value for the inferred admixture times is small, models using sample ages alone explain the data variance even less well at only 2.1%.

#### Steppe Yamnaya

The MesoNeo dataset provides more Yamnaya related individuals than were previously available, adding more data points for dating the admixture event between EHG and CHG groups. While this is a substantial boost in sample to size, there are still relatively few samples and therefore not enough power to detect any trends in admixture time with sample age or geography. This is compounded by the samples being from disparate geographic locations in Ukraine, Poland, Kazakhstan and across Russia.

Despite this, we obtained admixture time estimates from 16 of the 17 Yamnaya samples present in our MesoNeo subset (Fig 10). The genetic formation of the Yamnaya population is not well understood. Culturally they appear in the archaeological records 3300-2600 BC [32]. Most of our estimated admixture dates are 5000-6000 years before present, a millennium or so before their believed cultural formation. Three outlier samples have inferred admixture times of more than 7000 years ago. These all have large standard errors on the estimates and so are less reliable. However, even allowing for larger confidence intervals based on higher standard errors, their admixture times are placed well before 6,000 years before present.

**Fig 10.**
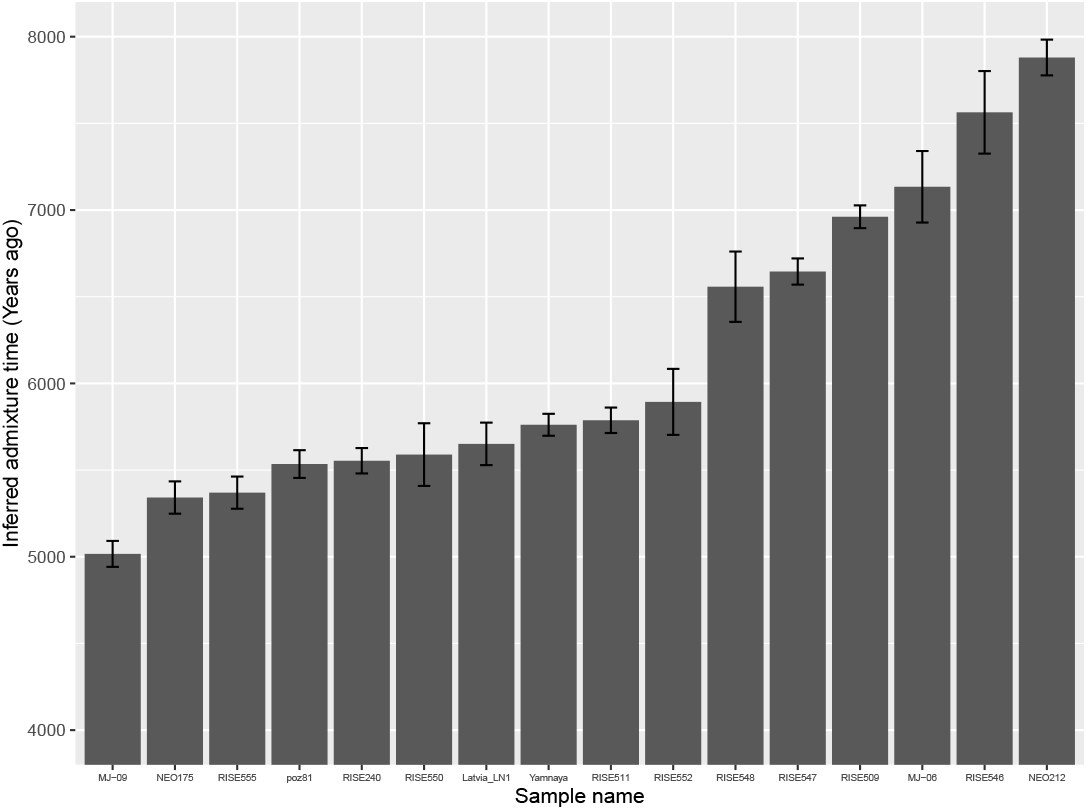
Inferred admixture times for Yamnaya samples. Barplot of the inferred admixture times for Yamnaya samples, with error bars showing the standard error around estimates.

These admixture dates are much older than the archaeological appearance of the Yamnaya and so, subject to technical artefacts they suggest extended prior genetic contact between EHG and CHG well before the emergence of the Yamnaya culture, potentially early as 7000 years ago.

## Discussion

Here we have presented a method for inferring local ancestry in ancient and modern genomes given an explicit model population history. The redefinition of local ancestry as a path, rather than a static identity, back in time through various populations is more appropriate to complex population histories. It is becoming more apparent, especially from ancient DNA analysis, that the histories of human populations across the globe are characterised by multiple population split, admixture, migration and isolation events and an appropriate analogy is a braided river rather than a tree. Hence, with the growing amount of ancient DNA data available, the path concept of ancestry is becoming more viable with and applicable to populations other than Europeans. We suggest that it is an important step to begin thinking of ancestry in this way rather than in terms of static identity. It is this reframing of the meaning of ancestry which separates our method from pre-existing local ancestry inference tools.

We have used a highly interpretable machine learning framework to perform a process that could be done manually. This helps to avoid the pitfalls of a ‘black box’ where it would be difficult to know which basic features are driving classification. Our method performs as well as a leading local ancestry inference tool, GNOMix, for populations that are close to the time of admixture and performs better for populations with more time since admixture.

The method has its foundations the ability of RELATE to correctly construct a tree sequence. In essence we are embedding a tree sequence within a population structure and thereby assuming that the trees fit within the model constraints and reflects the underlying population structure. There are two reasons why this may not be the case: Firstly, any model of the past demographic structure of a population will be inaccurate. In most cases the structure will be simplified to represent only major population events and ignore smaller scale migrations. For some populations, there may be no pre-existing knowledge of the population structure and results from PCA and ADMIXTURE analysis may be unclear. While we have shown in Supplementary note S3 that the neural network can generalise over misspecification in the model, but with too little understanding of the history we may not obtain results of much confidence or meaning. Secondly, RELATE may not infer the tree sequences correctly, despite the model representing the true structure well. We have attempted to compensate for biases in the RELATE inference by training the classifier with GNNs extracted from RELATE inferred trees paired with labels from the true simulated trees. Results on simulations indicate this is successful.

We applied our method to a large imputed dataset of ancient European and West Asian genomes and use a subsequent technique to infer the time since admixture of ancient individuals. Imputation borrows information from closely related, higher coverage individuals, making samples that coalescence most recently with each other, more similar to each other than their true sequences actually are. Intuitively this effect should help the topology building process of RELATE by helping identification of nearest neighbours. Conversely, there is no way for imputation to recover variants that are private to ancient groups and so not present in the reference panel which will hamper neighbour identification in ancient individuals. Overall, we suspect the affect of imputation on the accuracy of inferring tree topologies is minimal, however pervasive imputation errors are is topic of research [33] and further analysis is needed to evaluate how much these errors might be affecting RELATE inference.

Additionally, in the local ancestry painting phase errors will create switches of ancestry across homologous chromosomes. As a result, phase errors will cause ancestral tracts to appear smaller than in truth and therefore produce admixture times that are older than in truth. It is possible that our calculated admixtures times have overestimated absolute times as we have not accounted for phase error. A future strategy to implement that accounts for phase errors is to use both haplotypes from an admixed sample when recording the changes in ancestry at increasing genetic distance and not treat the two chromosomes from the same sample independently. Switches in ancestry across the homologous chromosomes will be accounted for by measuring over both haplotypes.

However, our conclusions drawn from the relative admixture times between samples are still valid, revealing spatiotemporal patterns of admixture of different populations in Europe. We showed how the inferred admixture date is superior for identifying these patterns than using the archaeological sample age alone. Results from Neolithic farmer genomes suggest that the movement of Anatolian farmers into Europe and their subsequent admixture with WHG was slow and can be explained by geography to a large extent. Admixture appears to occur first in Iberia and moves north west over time, potentially supporting a route of Anatolian farmers along the north Mediterranean coast into Europe. In contrast, results from Bronze Age genomes suggest that the movement of steppe Yamnaya into Europe was fast with admixture occurring soon after migration. We found that the genetic formation of the Yamnaya is 5000-6000 years before present, approximately a millennium before their believed cultural formation determined from archaeology which is consistent with other recent results [34]. We also show evidence of potential contact between the EHG and CHG in the eighth millennium before present.

It is important to note that admixture times are not always a proxy for movement of people. It is possible that the Anatolian Farmers moved at a faster pace into Europe, but the subsequent integration with the local hunter gatherer populations was a slower process that continued long after migrants arrived in an area. There is evidence of persistent un-admixed hunter gatherer pockets existing long after the arrival of Anatolian migrants to the same area which is seen in parts of the World today, such as the Hadza in East Africa.

For populations where the historical structure is well understood, the method can be adapted to these populations. For those where a demographic model from which to simulate is not available, one must be custom built, although the demographies of many species and human populations are now available in the stdpopsim catalogue (https://github.com/popsim-consortium/stdpopsim/blob/main/stdpopsim/catalog/HomSap/demographic_models.py).

## Supporting information

Code is available at the following link: https://github.com/AliPearson/AncestralPaths

