## Supplementary Information for "Local Ancestry Inference for Complex Population Histories"

S9 Summary of longitude and latitude linear models for 97 Bronze Age samples. 22

### S1 Excluding Bronze Age Anatolians

Our model of European population structure includes the Bronze Age Anatolians (BAA) from which paths 5 and 6 lead. Given these paths are not directly relevant to the history of present day Europeans, we tested how the accuracy of classification is altered when the BAA are removed from the GNN distributions and paths 5 and 6 are removed as labels. This leaves a four path model and a reduced GNN matrix size over which to train a neural network. We tested the accuracy of this classifier in each of the 4 path classes, averaged over 5 testing tree sequences, in the GBR population.

The overall accuracy across the four paths in the model with no BAA is 94.3% +/- 0.27%. This is significantly greater than the overall accuracy in the model containing BAA when taken over all six paths in that model (p-value = 1.473e-05). However when averaging over just the four non-BAA paths in the model containing BAA we obtain an accuracy of 94.6% + 0.46%, which is not significantly different to the classifier trained on the model excluding BAA (p-value = 0.0918).

This result can be seen in Figure S1 where the precision in each of the four non-baa paths in the model including BAA is very similar to those in the model excluding BAA.

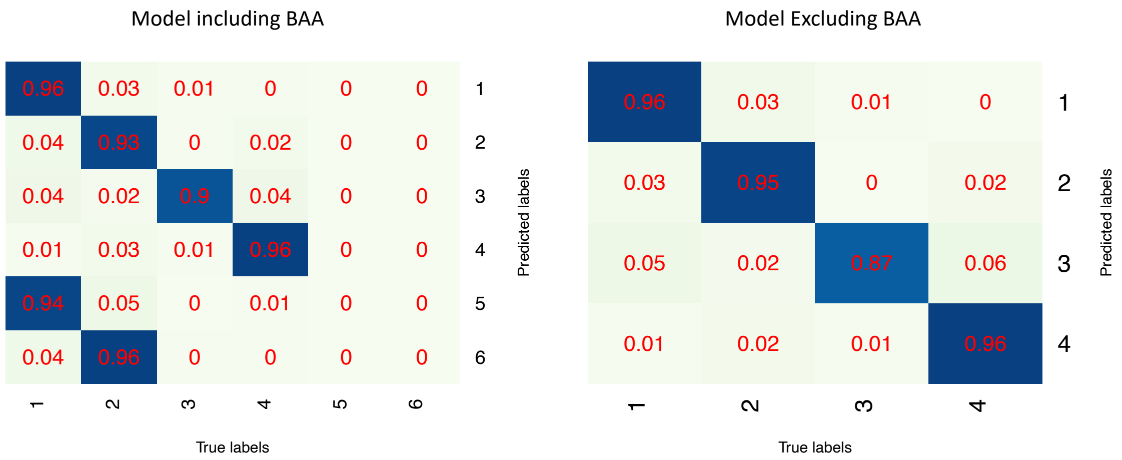

**Fig S1.** Confusion matrices showing the precision in each path tested over GBR samples in 5 testing tree sequences. The classifier trained on the model including BAA can predict one of six paths, while the classifier trained on the model excluding BAA can predict one of four paths.

This result demonstrates that accessory paths such as those leading from the BAA in our model of European population structure can be added with no detriment to precision of other paths. Although the overall accuracy across all paths is lower when the BAA are included, the accuracy is still over 90%.

### S2 Conditions affecting classifier performance

The "path ancestry" inference approach outlined above is applicable to many populations of humans and other species. To help decide what types of populations and time resolutions this method would be appropriate for we assessed its performance under a variety of demographic scenarios. we simulated data from a range of demography models, systematically varying features and parameters and we measured the overall classifier accuracy and precision in each class when tested on simulated data.

Unless otherwise specified, we simulated tree sequences of 20 Mbp in length with 1.25e-8 mutation rate and constant recombination rate of 1e-8. All admixture events involve equal proportions contributed by each participating population. The ordering of population split and admixture events is determined randomly by sampling from the active populations. The timing of split events is three generations after the previous split event while the timing of admixture events is fifteen generations following the previous admixture event. All populations have an effective population size of 10,000. Twenty five diploid samples were taken from all the 'path' populations that are present after all split events have occurred and also from all admixed populations. The sampling time for each population was sampled uniformly from within the time each population was active and all samples per population are taken at the same time. An example is shown in Figure S2.

Training GNNs and label pairs were extracted from all admixed populations. To gather the same number of pairs for each path in each admixed population from sufficiently spaced sites, we simulated five training tree sequences and five testing sequences. The largest source of variation is across tree sequences so testing was performed on each sequence separately and a mean and standard deviation across sequences was calculated. Two sample t-tests were then performed to assess if there was a significant difference in classifier accuracy when a parameter was changed. We then pooled the testing GNNs and applied the classifiers to produce confusion matrices and calculate precision values in each class.

### S2.1 Inference with varying path number and divergence time

We tested how the number of paths in the model and how much differentiation between the 'path' populations affected the classifier's ability. This was to explore 1. how complex the demographic history could be for the classifier to tease apart separate paths and 2. how applicable the method is to more finely structured populations with recent admixture compared to populations with deep structure.

We simulated demographic models that contained two, four, six and eight paths. All demographic models started with a single trunk population that through binary population splits, divides into several populations corresponding to the number of paths. These populations remain separate for around a specified number of generations (10, 50, 100, 500, 1500 or 3500) before admixing successively with each other until one population remains (Figure S2). The same random seed was used to simulate models of the same path number, so all models with the same path number but different divergence times had the same ordering of population splits and admixtures.

Figure S3 shows the accuracy and standard deviation for classifiers trained on all demographic combinations of path number and separation times. The accuracy decreases as the number of paths in the model increases and as the number of generations that all the paths are diverged decreases. All path numbers show a rapid increase in accuracy from 100 to 500 generations of divergence. The more generations that the 'path' populations are separate, the larger the allele frequency differences become between paths for pre-existing variants due to drift acting for a longer period of time. Additionally more mutations accumulate that differentiate paths while paths are separated. More mutations and greater allele frequency differences means that RELATE is better able to resolve the correct topologies, which results in the GNNs looking more consistent within each class.

More paths increases the number of classes that the classifier must differentiate between and therefore the opportunity to confuse between those classes. Models with more paths will require more time of divergence to achieve the same accuracies as models with fewer paths. Models with only two paths maintain accuracies of above 50% with as few as ten

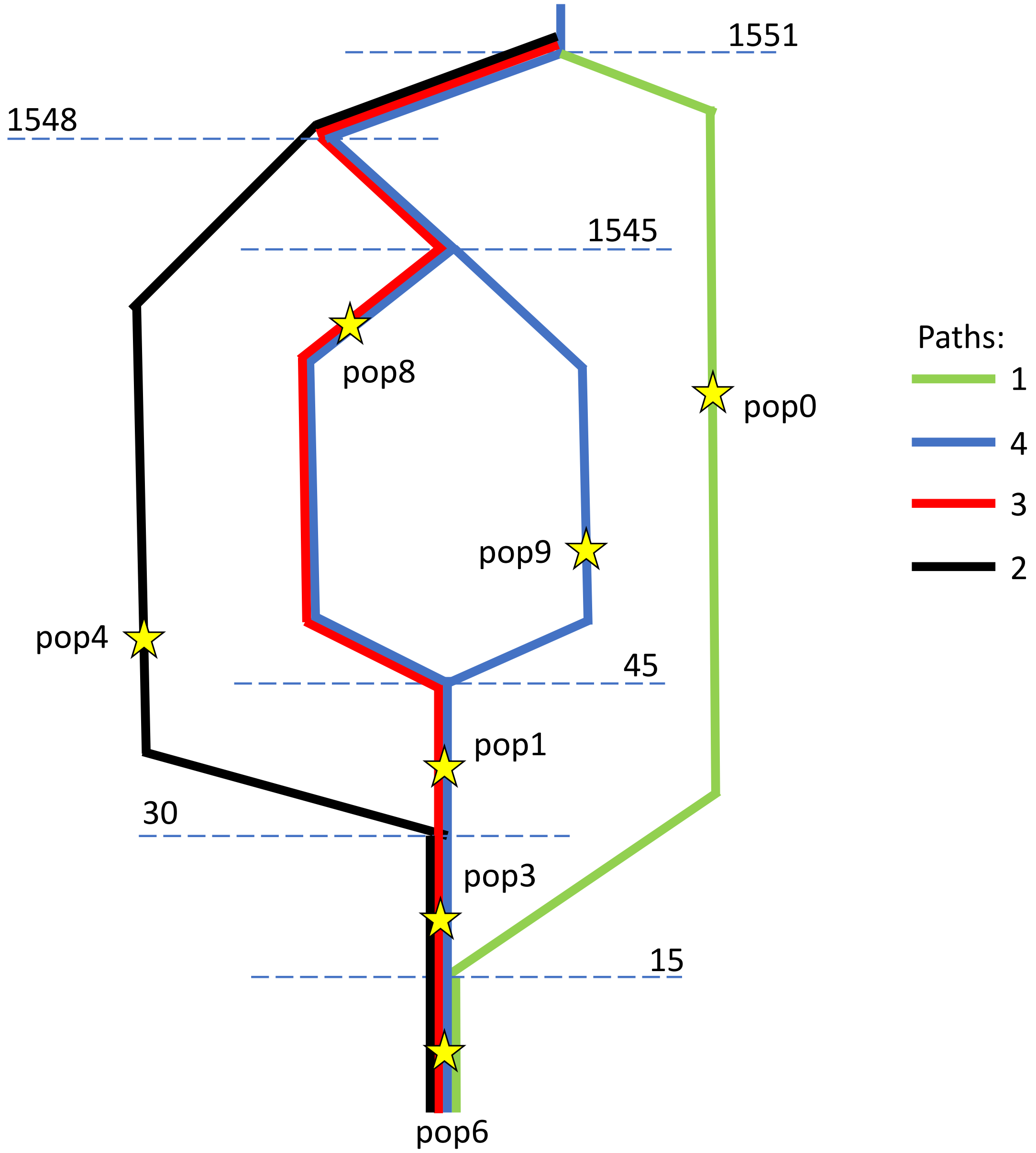

**Fig S2.** An example of a four path demography that was simulated for testing. Stars indicate approximately where in time 25 diploid samples are taken. The populations and paths are arbitrary numbered. Population split and admixture times are shown on the dotted lines in units of generations ago. This example has a ‘path’ divergence time of 1500 generations, the time period between the last population split and the first admixture event.

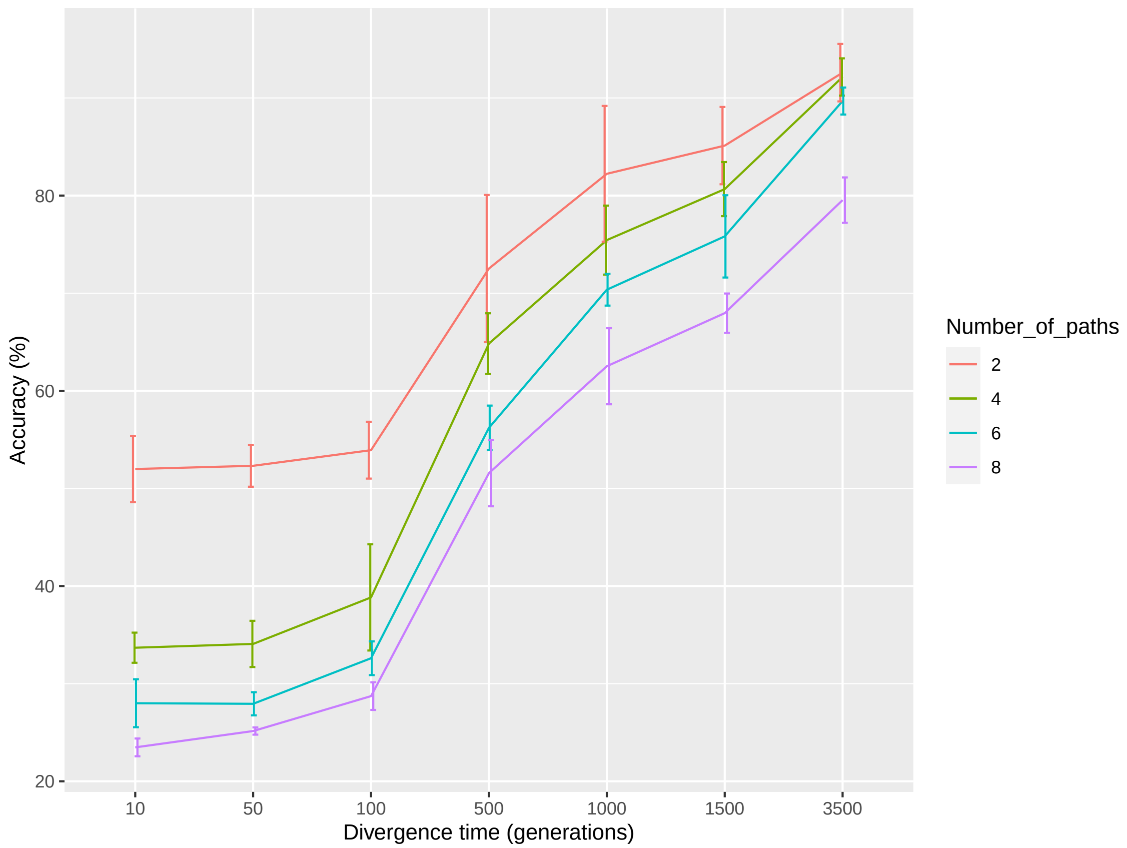

**Fig S3.** Plot showing the mean and standard deviation of accuracy of classifiers trained on models with different path numbers and 'path' population divergence times.

generations of separation time. By 3500 generations (∼100,000 years) of divergence, models containing two, four and six paths have accuracies above 90% and overlapping error bars, showing that with enough time for divergence the decrease in accuracy due to path number can be mitigated.

Overall, models with smaller path numbers are better suited when there is very fine scale, recent structure involving closely related populations. Likewise, when the populations are very diverged and deep structure is present, models containing more paths are viable.

### S2.2 Inference with imbalanced population size

Next, we tested how imbalance in population size on paths affects classification. We used a demographic model of four paths and 1500 generations of separation between 'path' populations. One 'path' population was chosen to have a population size of 50,000 from its emergence after a split to its disappearance after an admixture. All other population sizes were 10,000. A control set of tree sequences were simulated with the same population split and admixture events but with all population sizes set at 10,000.

Compared to the control demographic with an accuracy of 80.66% +/- 2.48%, the classifier trained on the imbalanced population size demographic demonstrates a significantly lower accuracy of 73.45% +/- 0.71% (two sample t-test p=0.0033). Comparing the confusion matrices (Figure S4), path 3, containing the differently sized population, exhibits the drop in precision and is mostly confused with path 4. This behaviour is not unexpected given paths 3 and 4 are sister paths, descending from a common population that split (Figure S2). A larger population size will reduce the number of coalescence events occurring around the time of higher population size compared to the other paths, pushing coalescences into older time periods before paths 3 and 4 separated. Many GNNs for path 3 will not have a coalescence event falling in the period of higher population size, making them look like path 4 or other GNNs and so are misclassified.

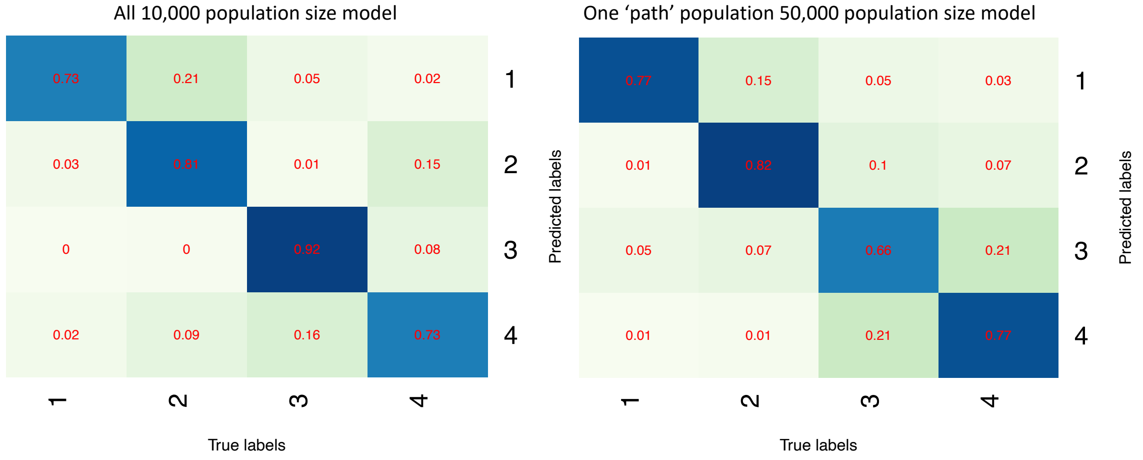

**Fig S4.** Confusion matrices of the model with all populations with size 10,000 compared to the model where one population has size 50,000.

### S2.3 Inference with imbalanced sampling

It is characteristic of ancient DNA datasets that some populations may only be represented by a few samples, while others many. Rather than subsetting some groups to match the sample size of the groups with the fewest samples, we tested if an imbalanced number of

**Table S1.** Table displaying the mean and standard deviation of accuracy of classifiers trained on models with variable number of samples taken from a population along path 1.

| Number of samples from path 1 population | Mean accuracy (%) | Accuracy standard deviation (%) |
|---|---|---|
| 5 | 80.22 | 4.48 |
| 25 | 80.66 | 2.77 |
| 50 | 82.49 | 2.65 |

samples taken from each group alters the classification of certain paths. we simulated from three demographics with divergence time of 1500 and four paths and trained a classifier to each. In the control demographic model, the sample size was 25 diploids for all 'path' and admixed populations. In the other demographic model, one 'path' population sample size was either reduced to 5 diploids or increased to 50 diploids. The same order and timing of split and admixture times were used, and all population sizes were 10,000. Path 1 contained the population with variable samples.

There was no significant difference in overall accuracy of the classifiers when tested pairwise with two sample t-tests (p-values = 0.318, 0.856, 0.364). Table S1 shows the accuracies for each model and the standard deviations. The model with 5 sampled diploids on path 1 has the highest standard deviation. This is likely because there are fewer examples of path 1 labels from those samples in the training data and so the training has not captured as much of the variance of path 1 GNNs as other classes when training the classifier. Testing that classifier, this translates to greater variance in the accuracy.

In the confusion matrices (Figure S5) path 1 has greater precision in the 50 diploid model than the other two models, which have comparable precision values in path 1. However, the overall mean accuracies are not significantly different suggesting there is no systematic bias due to imbalanced sample sizes, rather an increase in precision due to greater sample size on one path.

### S2.4 Inference and overall sample size

Lastly, we tested the effect of overall sample size on classification ability. We simulated from seven different demographic models with a divergence time of 1500 and four paths and trained a classifier to each. Each had a different number of diploids sampled evenly from the admixed and 'path' populations: 5, 10, 15, 20, 25, 30 or 50 diploids. The same order and timing of split and admixture times were used, and all population sizes were 10,000. The same number of training GNNs were used to train classifiers for each demography so as not to confound results with different amounts of training data.

Figure S6 shows how the mean accuracy increases as the number of diploids taken from all sampled populations increases. There is a rapid increase in accuracy between 5 and 20 diploids, after which the increase in accuracy with more samples is less. At 50 diploids the

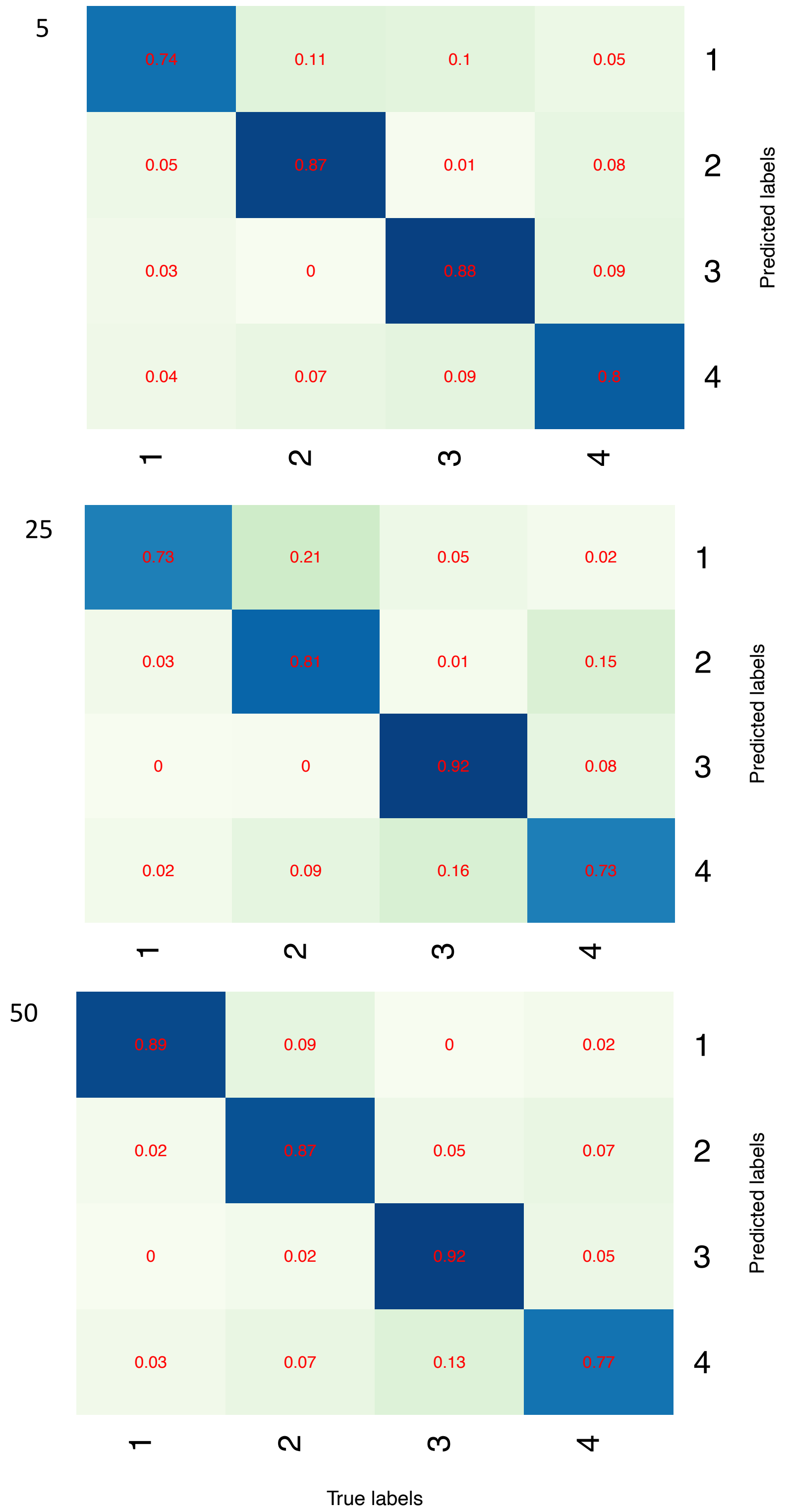

**Fig S5.** Confusion matrices of models with different number of diploids sampled from a path1 population.

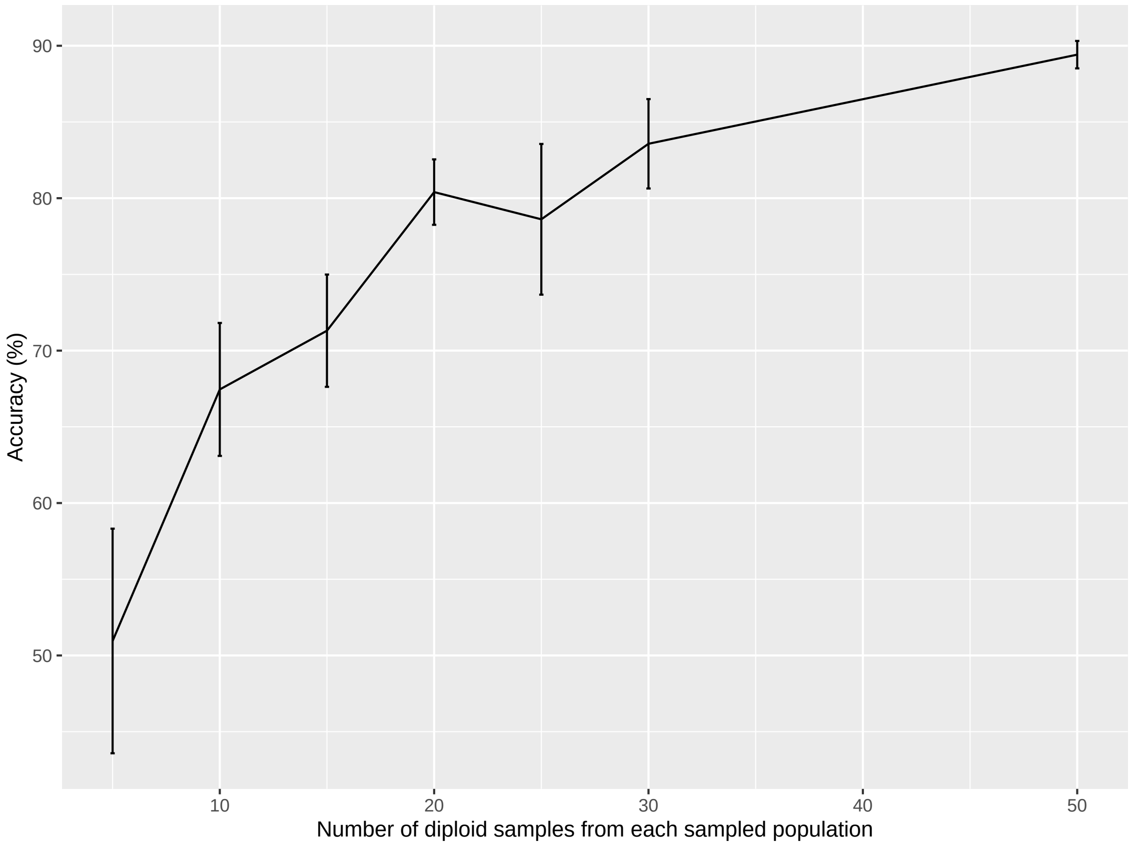

**Fig S6.** Change in accuracy as the number of diploids sampled from all admixed populations increases. Errors bars show the standard deviation over five testing tree sequences.

accuracy is around 90% and the standard deviation is low. Even with only 10 samples, with a 1500 generation diverged four path demography, the accuracies can be over 70%.

More samples means more variation in the training GNNs, given that we used the same number of training GNNs from each demography. This prevents overfitting of the neural network allowing for greater flexibility to novel testing GNNs. This results in greater accuracy and smaller standard deviations upon testing.

### S3 Testing model misspecification

The neural network for determining European local ancestry is trained on simulated data that we believe to match the real data well enough. However, the true history of any population is never known exactly and the following section explores how a classifier copes when it is trained on data that does not match the testing data in various ways. For all investigations we used demographies with four paths and 1500 generations of path separation time. Except where specified, simulations were carried out in the same way as described in Subsection S2.

For each parameter under scrutiny, we simulated two datasets with only the parameter under scrutiny differing between the two and train a classifier to each. Testing within the datasets was performed as in Subsection S2. When testing between datasets, the classifier of one was applied to each of the five testing tree sequences from the other dataset. This results in a mean accuracy and standard deviation for all four combinations of classifiers and testing data, to which we applied two sample t-tests to determine whether there is a significant difference in classifier accuracy when applied to testing data that does not match the training data. The two classifiers were then applied to pooled testing data in all four combinations to produce confusion matrices.

### S3.1 Inference with different population size

Using the same simulations and classifiers as those used for testing an imbalance in population size along paths, we tested the two classifiers trained on a 10k model and an imbalanced 50k model on GNNs extracted from the alternative demography. A difference in population size in a 'path' population will change the GNN structure, where a smaller population size will produce more coalescence events in the path population and a larger population size will conversely produce fewer coalescences. A difference in either direction in the GNN structure between training and testing data will result in the paths being less easy to recognise by the classifier. Predictably there is a significant decrease in accuracy when test data is classed by the classifier trained on different training data to when classed by the classifier trained on the corresponding training data (Table S2). Two sample t-tests produce p-values of 0.007 and 0.0067.

There is no significant decrease in accuracy when the 50k classifier is applied to 10k data compared to 50k data (p-value = 0.418). However there is a significant decrease in accuracy (p-value = 9.661e-05) when the 10k classifier is applied to 50k data compared to 10k data. An unexpected deficit of coalescences along a path is more confusing to a classifier than a surplus i.e the absence of a defining GNN is more detrimental than the presence of an extra GNN.

**Table S2.** Table displaying the mean and standard deviation of accuracy of classifiers trained on models with the same or different population size than the testing data, along path 3.

| Testing data / Classifier | 10k all | 50k imbalanced |
|---|---|---|
| 10k all | 80.66. +/- 2.77 | 68.95 +/- 2.12 |
| 50k imbalanced | 74.52 +/- 2.61 | 73.45 +/- 0.71 |

#### S3.2 Inference with different admixture times

To explore how the classifier generalises to data with different admixture times, we simulated from one demographic model with admixture events occurring every 15 generations back in time from the present day and another every 30 generations. So the latter did not result in a smaller time while the 'path' populations were diverged, which would confound the results, we increased the time of the most recent population split in that demographic model.

Table S3 shows that the accuracy for the model with 30 generations separating admixture events is not significantly different than the accuracy of the model with 15 generations, when tested on their corresponding testing data and trained classifier (p-value = 0.976). When we swap the classifiers, to test the 30 generation separated testing data with a 15 generation separated classifier and vice versa, there is no significant change in accuracy in either case compared to testing corresponding data and classifiers (p-values = 0.977, 0.716). The classifiers are able to generalise and compensate for the difference in admixture times. This indicates that, despite providing the node ages in the GNNs, the classifiers are largely using the topology of trees and not the coalescence times.

**Table S3.** Table displaying the mean and standard deviation of accuracy of classifiers trained on models with the same or different admixture times than the testing data.

| Classifier \ Testing data | 30 gen. admixture | 15 gen. admixture |
|---|---|---|
| 30 gen. admixture | 80.58 +/- 4.81 | 81.27 +/- 2.33 |
| 15 gen. admixture | 80.49 +/- 4.65 | 80.66 +/- 2.77 |

#### S3.3 Inference with different admixture fractions

We next investigated how different admixture fractions would change the classification accuracy. We simulated a dataset with all admixture fractions 50/50 and another with all 25/75. There is no significant change in accuracy when the 50/50 classifier is applied to 25/75 testing data compared to testing the corresponding 25/75 data and classifier (p-value = 0.393). Neither is there a significant change in accuracy when the 25/75 classifier is applied to 50/50 data compared to testing the corresponding 50/50 data and classifier (p-value = 0.589). This suggests that classifiers are able to generalise when the admixture fractions are misspecified.

#### S3.4 Inference when samples are drifted from ancestral populations

For all models we have been simulating samples that are taken directly from the simulated populations. The ancient samples we have in the MesoNeo dataset are unlikely to be individuals from the ancestral populations involved in the admixture events themselves, but

**Table S4.** Table displaying the mean and standard deviation of accuracy of classifiers trained on models with the same or different admixture fractions than the testing data.

| Classifier \ Testing data | 50/50 | 25/75 |
|---|---|---|
| 50/50 | 80.66 +/- 2.77 | 85.53 +/- 1.95 |
| 25/75 | 79.58 +/- 3.27 | 86.82 +/- 2.51 |

instead more or less closely related to them. To investigate the effect of drift between the true ancestral admixing populations and sampled populations, we simulated population splits so that the ancient samples were taken from 'hanging branches', slightly diverged from the admixing lineages. A separation time of ten generations from the true ancestral populations simulates approximately 300 years of drift. We trained classifiers using a demographic model where the ancient samples are taken directly from the ancestral admixing populations and tested its performance on GNNs from the drifted model.

**Table S5.** Table displaying the mean and standard deviation of accuracy of classifiers trained on models with ancient samples that were drifted or not from the true ancestral populations.

| Classifier \ Testing data | Drifted | Not drifted |
|---|---|---|
| Drifted | 83.28 +/- 1.44 | 84.60 +/- 0.78 |
| Not drifted | 81.51 +/- 3.32 | 82.63 +/- 3.79 |

Table S5 shows the accuracy results when classifiers are testing on different testing data. There is no significant change in accuracy between any two pairs of testing data and classifier combinations demonstrating that classifiers are able to generalise between drifted and directly sampled models (p-values = 0.106, 0.314, 0.634, 0.320, 0.733, 0.122).

### S3.5 Inference for additional samples

The number of ancient samples that are available continues to increase and so more samples relevant to population histories can be incorporated into datasets. Likewise, samples may be removed from datasets after more stringent filtering. To test whether classifiers can generalise to more or fewer samples, we applied a classifier trained on a simulation with 20 diploids taken from all sampled populations to testing data from a simulation with a different number of diploids per sampled population (5, 10, 15, 25, 30, 50). GNNs were extracted using all samples in the demography, as if testing a corresponding classifier.

Figure S7 shows that there is no significant difference when using the classifier trained on 20 diploids compared to the corresponding classifier for testing data containing 5, 10, 15, 25 ad 30 diploids (p-value¿0.05). For data containing 50 diploids there is a marginally significant decrease in accuracy using the 20 diploid classifier.

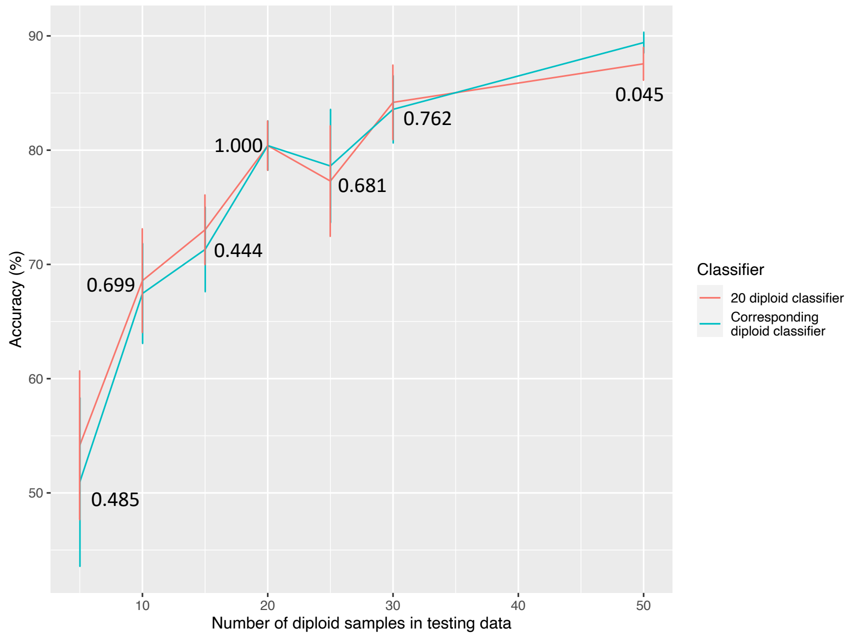

**Fig S7.** Change in accuracy as the number of diploids sampled from all sampled populations increases using a classifier trained on 20 diploids vs the corresponding classifier trained on the same number of diploids as the testing data. Errors bars show the standard deviation over five testing tree sequences. Numbers show the two sample T-test p-values comparing the two classifiers.

Classifiers were able to generalise to fewer samples than in the training data. When the number of samples is much larger than simulated in the training data, the accuracy begins to decrease. This decrease is only to a small extent even for more than twice the number of samples demonstrating that classifiers are flexible to differences in sample size.

### S3.6 Inference of ghost lineages

Genetic evidence has revealed ghost populations in many species, including humans. This is when a population is inferred to have existed but is not sampled with DNA or in the fossil record. To test whether our method could be used in a case involving a ghost population, we applied a classifier trained on a simulation where one 'path population' of the four paths present was not sampled.

We tested two scenarios from Figure S2 : One where the samples from population 0 were removed meaning path 1 contained the ghost lineage; and one where samples from population 9 were removed meaning path 4 contained the ghost lineage.

Figure S8 shows that the precision in all paths is not decreased when path 4 contains the ghost lineage and the precision of the classifier to predict path 4 actually increases.

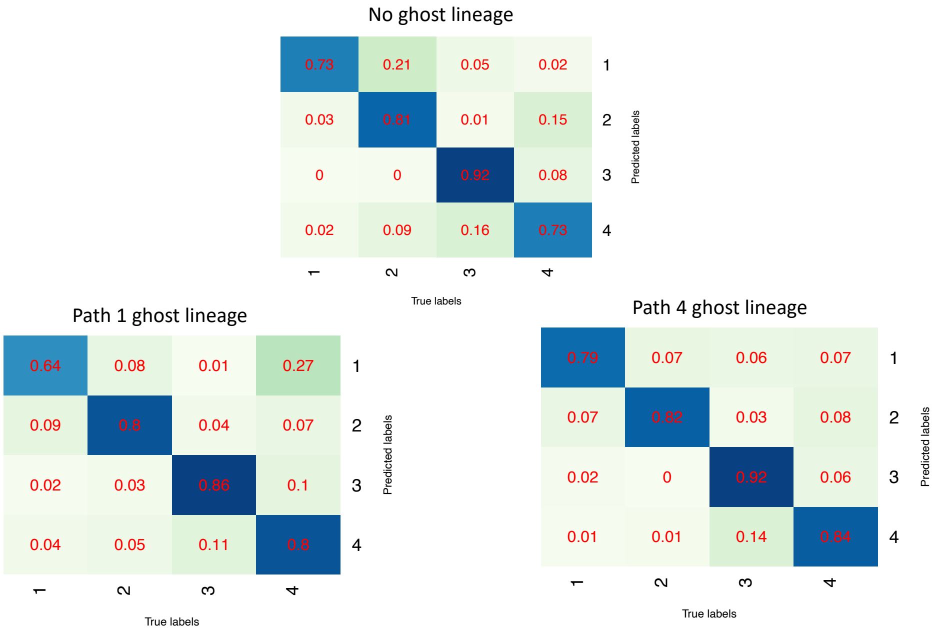

**Fig S8.** Confusion matrices showing the precision of classifiers in each class when various populations where removed from training and testing data to mimic ghost lineages.

It appears that the classifier receives enough information from other populations present along path 4 to continue to identify path 4 with high precision.

When path 1 contains the ghost lineages, the precision of the classifier is decreased in path 1. This is likely because there are no other populations along path 1 that are included in the GNNs, meaning that when population 0 is removed it becomes harder to identify the path.

Overall, the accuracy remains high for identifying ghost lineages. The precision maintained depends on whether there are other sampled populations involved that can inform the classifier of path identity. The ability to identify ghost lineages, even with little to no other populations present along the lineages highlights the advantage of a path structure compared to a frame work of single population identity that is used by other local ancestry inference tools.

### S4 A method to estimate time since admixture

We devised an LD-based method to infer the time since admixture of samples from our painted chromosomes. LD-based methods are more robust to noisy ancestry inference and therefore are appropriate to our data. Segments are painted by path, so admixture corresponds to when two paths join in a population history. This means that tracts made up of two or more paths combined can be used to date multiple admixture events, where one event that joins multiple paths follows other events that join two or more paths. For example, the joining of paths 1, 2, 3 and 4 in the Bronze Age admixture event in Figure **??** is preceded by paths 1 and 3 joining in the Neolithic farmers and paths 2 and 4 joining in the Yamnaya.

With empirical sampling we plot the probability of being in the same path as a function of genetic distance and fit an exponential decay curve to the distribution. The parameters of the exponential decay correspond to the values of interest, time since admixture and admixture fraction. This can be done for each sample chromosome individually and the results across the whole genome for each sample combined to give an admixture time estimate and admixture fraction for each individual.

The sampling process is as follows:

1. Sample a starting position uniformly from between 0 and 0.5cM from one end of the chromosome.
2. Record the starting path.
3. Move 1cM towards the end of the chromosome.
4. Record the path 1cM away.
5. Now move to a new starting position 1cM away from the previous starting position.
6. Repeat steps 2-5 until the end of the chromosome.

Repeat the above steps for testing distances 1-50cM in step 3 and 4. Over all autosomes, we can then calculate the probability of being in path y given starting in path x, $d$ cM away, for all path combinations of x and y, including x=y. The shortest distance is 1cM, as in ROLLOFF, to avoid the effect of background LD.

The probability when x=y decays exponentially with genetic distance and the rate of decay depends on the time since admixture. Parameters are determined by nonlinear least squares regression of the data to the formula

$$P(\text{same path}) = \alpha e^{(-\beta d)} + \theta \tag{1}$$

The probability will asymptote to the probability of being in path $x$ across the whole chromosome, which intuitively is the admixture fraction of path x, therefore the admixture fraction is given by $\theta$. The $\beta$ parameter represents the time since admixture in generations (Figure **??**).

The process is performed over all autosomes for each individual genome and homologous chromosomes are treated as independent. For admixture events between two populations, each individual has two estimates of the admixture age, one calculated per ancestral path. We combined the estimates of the two in a way that minimises the standard error to give a weighted average value of time since admixture for each individual. The time of admixture in generations ago from present-day can therefore be calculated as the sum of the sample age in generations ago and the estimated time since admixture for that sample.

### S4.1 Performance on simulated data

To test the performance of this method, we simulated data from the model of European population structure. We executed the analysis on the admixed populations to infer time since admixture and admixture fractions of paths.

We dated the admixture times of all individuals by extracting the $\beta$ parameter from fitting exponential decay curves as described above. Counts for all distances were taken across all nine tree sequences. The probability of remaining in the same path was therefore calculated using the total counts from across 18 independent sequences per individual. The method was applied on both the simulated painted tree sequences and on tree sequences that were inferred by RELATE from the simulated data and painted using a classifier. The results are shown in Figure S9 and Tables S6 and S7.

**Table S6.** Table of results of admixture time analysis performed on simulated painted tree sequences. For three admixed populations the mean time of admixture across all simulated samples and the standard deviation of estimates around the mean is shown. All values are in units of generations ago.

| Population | Simulated time of admixture | Mean time of admixture | Standard deviation (+/-) |
|---|---|---|---|
| Bronze Age | 166 | 166.03 | 3.86 |
| Neolithic farmers | 259 | 258.54 | 12.96 |
| Yamnaya | 177 | 177.74 | 1.66 |

In Figure S9, as the sample ages become more recent and further from the admixture time, the variance of estimates around the true value increases for both simulated and RELATE inferred results. Likewise, Figure S10 shows how the standard error of estimates increases as the sample ages get further from the time of admixture. More generations since admixture mean that more recombination events have occurred resulting in shorter ancestry tracts. Shorter tracts make it harder to differentiate between background LD inherited from the parent populations from the relevant admixture LD. Therefore the standard error for the $\beta$ parameters when fitting exponential decay curves is greater and the variance in $\beta$ parameters estimates is greater. This is the case for both simulated data and RELATE inferred data showing there is a limit to the length of admixture tracts, even in data with no noise, for inferring time since admixture accurately.

Noise appears as short tracts of misclassified correlated trees in RELATE inferred data,

**Table S7.** Table of results of admixture time analysis performed on tree sequences that were inferred by RELATE from the simulated data and painted using a classifier. For three admixed populations, the mean time of admixture estimate across all simulated samples and the standard deviation of estimates around the mean is shown. All value are in units of generations ago.

| Population | Simulated time of admixture | Mean time of admixture | Standard deviation (+/-) |
|---|---|---|---|
| Bronze Age | 166 | 166.45 | 4.56 |
| Neolithic farmers | 259 | 257.28 | 16.67 |
| Yamnaya | 177 | 177.24 | 2.00 |

which starts to become difficult to differentiate from admixture LD when the admixture tracts become comparable in length. The standard error values for the analysis of tree sequences inferred by RELATE up to 100 generations since admixture are very similar to those from the analysis of the simulated data (Figure S10). This suggests that up to approximately 100 generations since admixture the method is robust to noise introduced by classification error in the painted chromosomes. Similar results can be seen in Figure S9, as the sample ages become more recent and further from the admixture time, the deviation of estimates from the true value increases in both sets of data but to a greater extent in RELATE inferred data. Likewise, the overall standard deviation of the mean time since admixture estimate across all samples is slightly larger in the RELATE inferred data compared to the simulated data for all populations (Tables S6 and S7).

In results of analysis of the simulated data, the Bronze Age mean admixture time predicted across all samples matches the true value of 166 (Table S6) and a small standard deviation of estimates of +/- 3.86 generations. Neolithic Farmers display a mean admixture time slightly under the true value of 259 and a larger standard deviation of estimates of 12.96 +/- generations. The slightly poorer ability to estimate Neolithic farmer admixture time is because some samples have ages that are further from admixture compared to the Bronze Age population, resulting in the mean being slightly under the true admixture time and a greater standard deviation. The error may also be increased by the ancestry proportions of 75/25 Anatolian and WHG respectively, as the exponential decays are harder to fit when the difference between the starting value, near 1, and the asymptotic value of 0.75 is not as large. Yamnaya samples are a maximum of 17 generations from the true admixture time of 177. The mean predicted admixture date matches that simulated and there is a small standard deviation of estimates of +/- 1.66 generations.

Results of analysis of RELATE inferred data were very similar to that of the simulated data (Table S7). The Neolithic farmer's mean $\beta$ estimate is slightly further from the truth in the RELATE inferred data analysis than in the simulated data analysis due the same

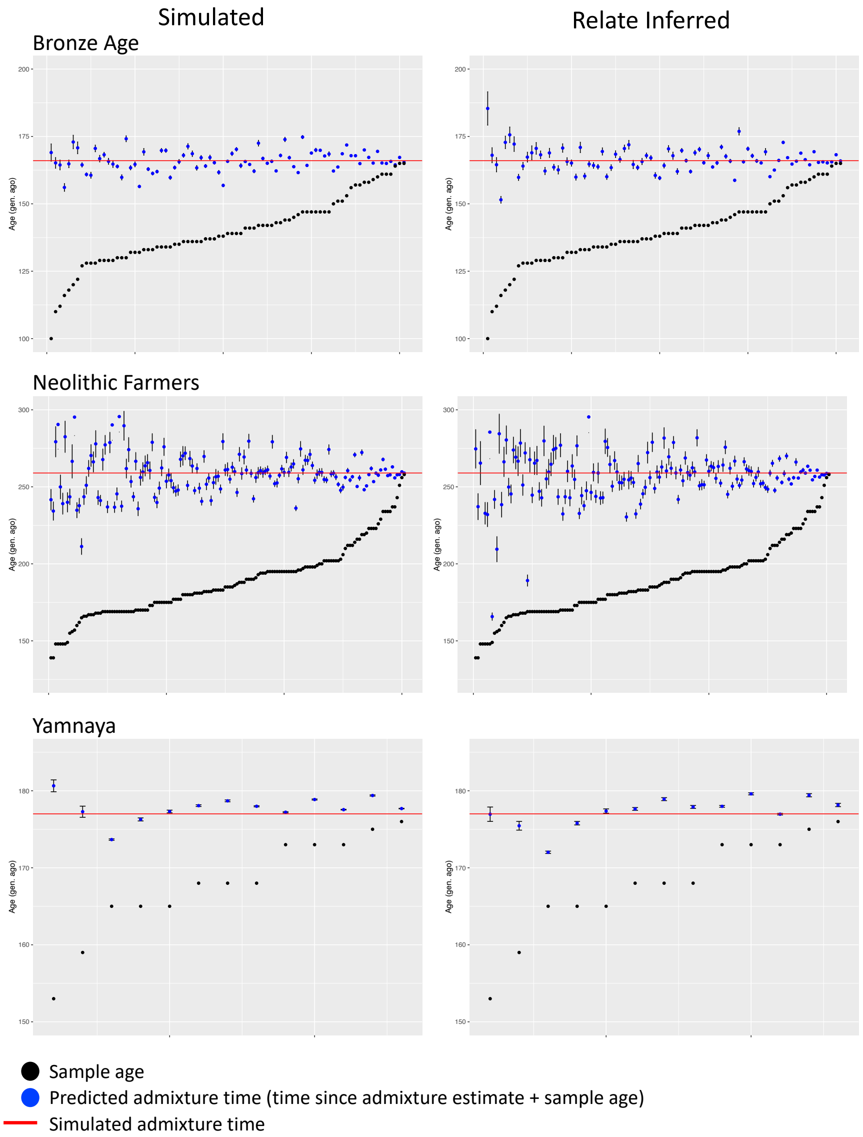

**Fig S9.** Plots showing the predicted admixture time with standard error bars and the sample age for three simulated admixed populations. The time since admixture from each painted individual was calculated from the $\beta$ parameter described in Section **??** and the time of admixture plotted from the sum of the time since admixture and sample age in generations ago.

reasons explained above; more samples with a greater time since admixture, above 100 generations, meaning noise introduced by classification has more of an effect of decreasing

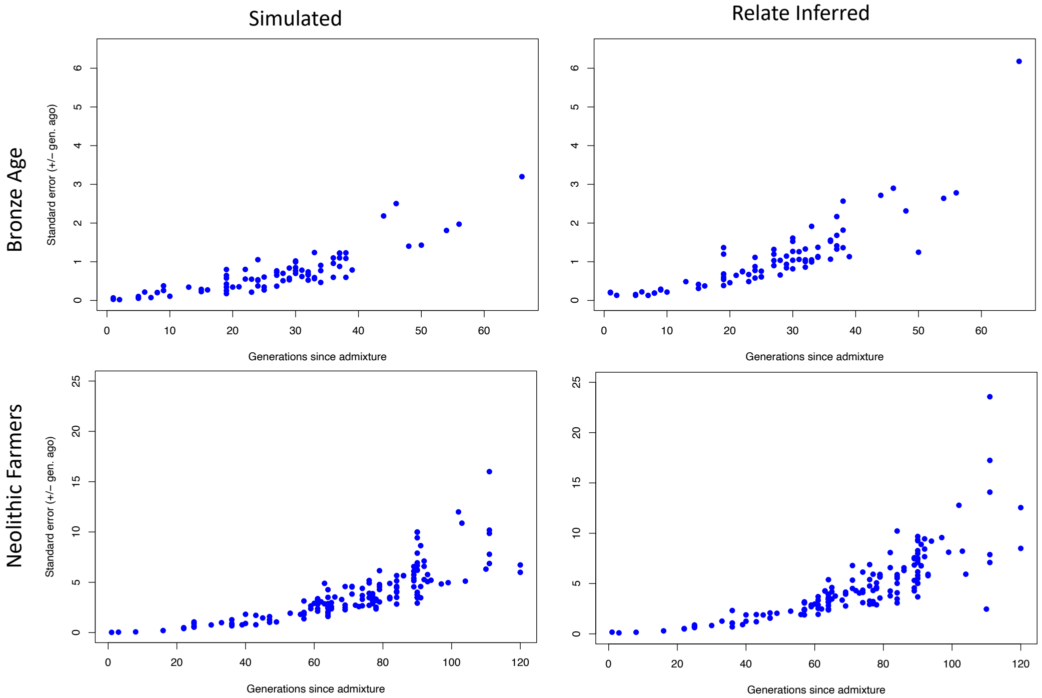

**Fig S10.** Plots showing the decrease in standard error of admixture time estimates as sample age increases and gets closer to the simulated time of admixture for Bronze Age and Neolithic farmer populations. The left column shows the results for admixture time analysis on the simulated tree sequences and the right shows the results for tree sequences inferred by RELATE from the same simulated data and painted using a classifier.

the accuracy of the $\beta$ estimates.

For the present-day samples that are all 166 generations since admixture, in both simulated and RELATE inferred tree sequences, the admixture tracts have become too short to produce a reliable estimate of time.

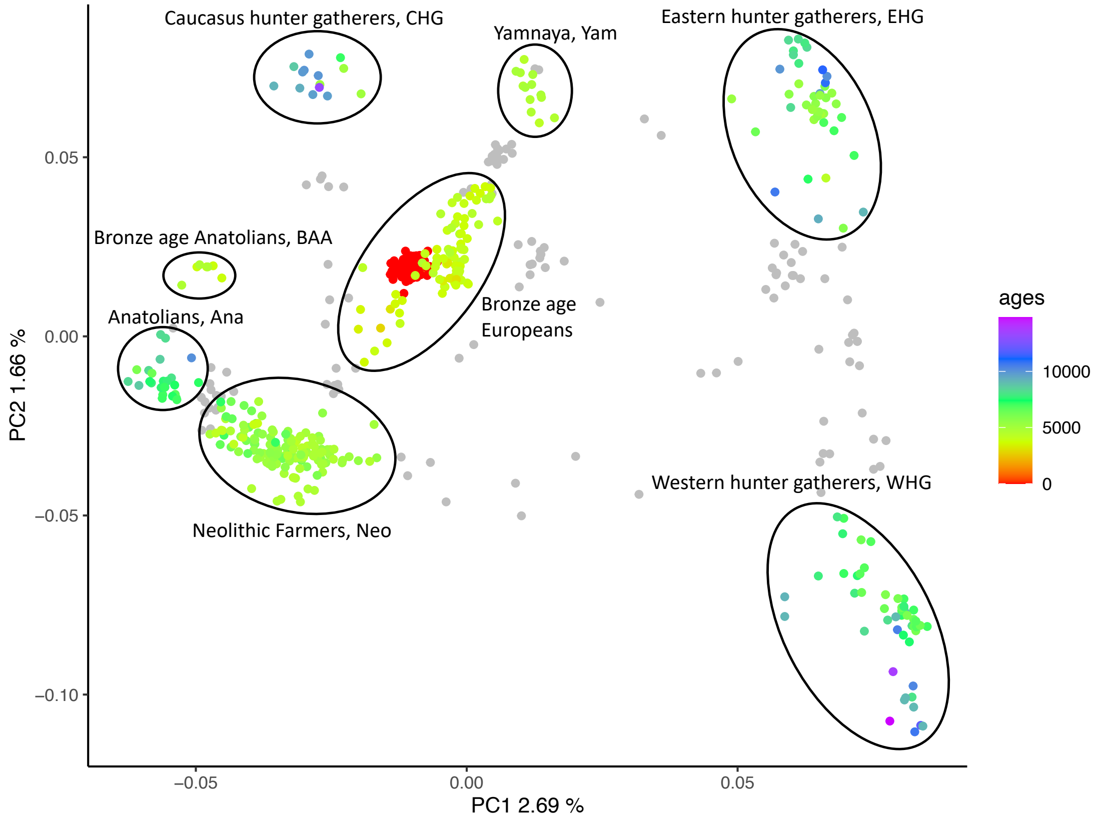

**Fig S11.** Subset of MesoNeo genomes, plotted by their first two principal components. Samples that are diagnostic of each ancient European group are coloured by their radiocarbon age in years ago. The samples that fall in between circled samples and were removed from further analysis, coloured in grey.

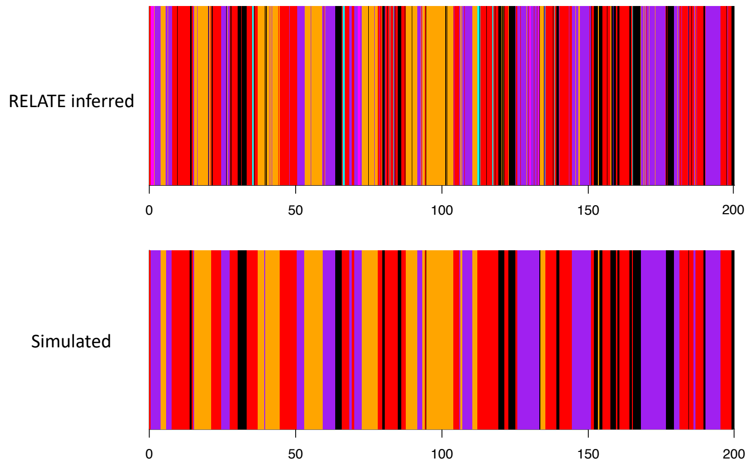

**Fig S12.** Example simulated painted haploid chromosomes. Painted haploid chromosomes from a Bronze Age individual . The top chromosome shows the true simulated painting and the bottom chromosome shows the corresponding RELATE inferred chromosome, painted by classification.

**Table S8.** Summary of longitude and latitude linear models for 173 Neolithic farmers samples.

| **Coefficients** | **Estimate** | **Std. Error** | **P-value** |
|---|---|---|---|
| Inferred admixture time $\sim$ longitude + latitude | | | |
| Intercept | 9092.66 | 450.59 | < 2e-16 |
| Longitude | -25.66 | 4.87 | 4.08e-07 |
| Latitude | -42.74 | 8.98 | 4.18e-06 |
| $R^2$-adjusted = 0.234 | | Model p-value = 5.947e-11 | |
| Sample age $\sim$ longitude + latitude | | | |
| Intercept | 7394.92 | 488.34 | < 2e-16 |
| Longitude | -11.70 | 5.65 | 0.0396 |
| latitude | -35.25 | 9.89 | 0.000471 |
| $R^2$-adjusted = 0.084 | | Model p-value = 0.0013 | |

**Table S9.** Summary of longitude and latitude linear models for 97 Bronze Age samples.

| **Coefficients** | **Estimate** | **Std. Error** | **P-value** |
|---|---|---|---|
| Inferred admixture time $\sim$ longitude + latitude | | | |
| Intercept | 3894.05 | 443.23 | $7.57e-14$ |
| Longitude | 2.98 | 5.77 | 0.607 |
| latitude | 16.15 | 9.08 | 0.0785 |
